# Optimization of the *TeraTox* assay for preclinical teratogenicity assessment

**DOI:** 10.1101/2021.07.06.451364

**Authors:** Jaklin Manuela, Zhang Jitao David, Schäfer Nicole, Clemann Nicole, Barrow Paul, Küng Erich, Sach-Peltason Lisa, McGinnis Claudia, Leist Marcel, Kustermann Stefan

## Abstract

Current animal-free methods to assess teratogenicity of drugs under development still deliver high numbers of false negatives, and more sensitive approaches of toxicity prediction are required. To address this issue, we characterized the *TeraTox* test, a newly developed multi-lineage differentiation assay for human teratogenicity prediction using 3D human induced pluripotent stem cells. *TeraTox* produces as primary output concentration-dependent data sets for each test compound on cytotoxicity and altered gene expression. These data are then fed into a prediction model based on an interpretable machine-learning approach. The final information obtained relates to the concentration-dependent human teratogenicity potential of drug candidates. We applied *TeraTox* to profile 33 approved pharmaceuticals and 12 proprietary drug candidates with known *in vivo* data. This way, it was possible to relate the test predictions to known human or animal toxicity. The *TeraTox* had an accuracy of 69% (specificity: 53%, sensitivity: 79%). It clearly performed better than two quantitative structure-activity relationship (QSAR) models and it had a higher sensitivity than the murine embryonic stem cell test (mEST) run in the same laboratory. By combining *TeraTox* and mEST data, the overall prediction accuracy was further improved. The knowledge on the pattern of altered gene expression may provide additional value in grouping toxicologically similar compounds and possibly deducing common modes of action. The assay will thus be a helpful additional tool in drug discovery, and the dataset provided here will be a valuable resource for the field of stem cell-based drug profiling.

## 1. Introduction

To assess the teratogenic potential of drug candidates, pharmaceutical companies are currently obliged to perform embryo-fetal-development studies (EFD studies) in at least one rodent and one non-rodent species (Beck et al. 1995; ICH 2020). There is an urgent need to develop alternative, animal-free assays for early assessment of teratogenicity. The use of humanized *in vitro* assays could potentially better mimic human physiology, reduce animal use and lower the cost of drug development by filtering out potential teratogens early (Barrow 2016; ICH 2020; Lenz and Knapp 1962).

Animal-free approaches based on *in silico* prediction or *in vitro* assays have been widely adopted. For instance, the CAESAR model (Cassano et al., 2010) and the P&G model (Wu et al., 2013) are two qualitative structure-activity relationship (QSAR) models for reproductive and developmental toxicity prediction. Well-established *in vitro* models for the detection of teratogenicity include the *DevTox*^*qp*^ assay from Stemina and the mouse embryonic stem cell test (mEST). *DevTox*^*qp*^ uses human induced pluripotent stem cells (hiPSC) to predict teratogenicity based on the ratio of ornithine and cysteine in medium supernatants (Adler et al. 2008; Augustyniak et al. 2019; Burridge et al. 2011; Dreser et al. 2020; Palmer et al. 2017; Palmer et al. 2013; Shinde et al. 2015; Worley et al. 2018).

The mEST assay uses the beating of stem cell-derived cardiomyocytes as a functional readout for teratogenicity prediction (Genschow et al. 2000; Genschow et al. 2004; Scholz et al. 1999a; Scholz et al. 1999b; Whitlow et al. 2007).

Despite their wide adoption and relatively good performance, all these methods share major limitations. *In silico* models fail to consider the complexity and adaptiveness of biological systems. Also, their performance *a priori* for new chemical spaces that were not used for training the model is uncertain. On the other hand, existing *in vitro* models, which rely on a single readout for prediction, have limited capability to probe the complex biological processes underlying drug-induced teratogenicity.

To overcome these limitations, we developed a new, humanized *in vitro* teratogenicity assay for the pre-selection of pharmaceutical candidates. The new assay, which we call *TeraTox*, uses ethically non-restricted hiPSC derived embryoid bodies (EBs) that differentiate spontaneously into all three germ layers, with expression of representative early developmental markers of each layer. The new assay gave promising results in a preliminary evaluation (Jaklin et al. 2020). This paper describes the subsequent characterization and critical assessment of the assay, including a detailed predictive algorithm. A panel of 45 pharmaceuticals with evidence of human teratogenicity profiles based on FDA classification was tested using a six-point concentration-response generating the largest dataset of pharmaceuticals so far in a single study about *in vitro* modeling of teratogenicity with reference to clinical or animal data. We adapted an amplicon-based RNA sequencing technique (a technology known as *Molecular Phenotyping*) to quantify the expression of germ-layer genes as well as pathway reporter genes. We benchmarked a variety of architectures of machine-learning models and identified the best-performing predictive model using factor analysis and random-forest regression. *TeraTox* performed favorably compared with other models, *i.e.* the prediction accuracy was comparable with that of both Stemina *DevTox*^*qp*^ and mEST assays (higher sensitivity, lower specificity) and higher than that of two QSAR models (higher sensitivity and specificity). More importantly, *TeraTox* offers insights into a multitude of changes caused by the compounds on gene, pathway and germ-layer levels, some of which corroborated their teratogenicity potential.

## 2. Material and Methods

### 2.1. Human iPSC derived *TeraTox* Assay

The *TeraTox* assay is built upon a commercially available hiPSC line (Gibco, A18945), which has indistinguishable gene expression profiles compared with embryonic stem cells (Burridge et al. 2011; Quintanilla et al. 2014). The cells form 3D EBs and undergo multi-lineage differentiation into all three germ layers (Jaklin et al. 2020). Prior to the assay, the hiPSC were tested with the TaqMan ScoreCard assay (Thermo Fisher) to confirm sufficient levels of pluripotency (Tsankov et al. 2015a). The EBs were spontaneously differentiated and treated with each reference substance over a time course of seven days in Elplasia 96w micro-well plates (Corning, 4442) using the ViaFlo 96 automated microplate pipetting device (Integra) for liquid handling. Compounds were applied to the EBs on day 0, day 3, and day 5 at six concentrations, together with EB medium and 0.25% DMSO solvent controls as the negative reference. To test for batch-to-batch variation, we included several positive reference compounds in multiple runs (e.g., hydroxyurea, valproic acid, SB431542, etc.). Cell viability was determined prior to gene expression studies on day 7 by measuring ATP release in supernatants with the CellTiter-Glo 3D assay (Promega, G9681) according to the manufacturer’s protocol to pre-specify appropriate testing ranges. Concentrations that showed less than 80% cell viability were excluded from the subsequent gene expression studies. CellTiter-Glo™ reagent (100 μl) was added and incubated for 5 min on a shaker to lyse the EBs. The plates were kept for an additional 25 min in the dark at room temperature for binding of the released ATP to the luminescent dye. ATP release in supernatants was measured with the spectrophotometer (Biotek, Vermont, USA). All cell culture media and reagents were obtained from Gibco (Thermo Fisher) unless otherwise specified. The overall cell culture and cytotoxicity protocols have been described previously in detail by Jaklin *et al.*, 2020.

Targeted gene expression profiling was performed in biological duplicates at six sub-cytotoxic concentrations using the molecular phenotyping platform described previously (Drawnel et al. 2017; Zhang et al. 2015; Zhang et al. 2014). The resulting 1,055 samples of differentiated EBs were lysed after 7 days in 350 μl MagNA Pure LC RNA Buffer (Roche Diagnostics) and purified using an automated MagNA Pure 96 system (Roche Diagnostics). The total RNA was quantified using the Qubit RNA Assay Kit (Thermo Fisher) on the Fluorometer Glomax (Promega). Total RNA, with a maximum of 10 ng from each biological replicate, was reverse transcribed to cDNA using Superscript IV Vilo (Thermo Fisher). Libraries were generated with the AmpliSeq Library Plus Kit (Illumina) according to the reference guide. Pipetting steps for target amplification, primer digestion, and adapter ligation were done with a miniature mosquito automatic pipettor (SPT Labtech). For the purifications before and after final library amplification, solid-phase reversible immobilization magnetic bead purification (Clean NGS, LABGENE Scientific SA) was performed on a multidrop automated pipetting station (Thermo Fisher).

We measured both amplicon sizes and cDNA concentrations using an Agilent High Sensitivity DNA Kit (Agilent Technologies) according to the manufacturer’s recommendation. Prior to sequencing, cDNA contents of the samples were normalized and pooled to 2 nM final concentration on a Biomek FXP workstation. The libraries were sequenced on the NovaSeq 6000 Instrument (Illumina) using sequencing-by-synthesis technology. All 75 cycles ended up with a minimum of 2 Mio sequencing reads per sample for analysis. We used molecular phenotyping with 1,215 detectable pathway reporter genes, including a subset of 87 early developmental markers (germ-layer genes, Suppl. Tab. S4), and genes representative of toxicological pathways to identify differentially expressed genes induced at pre-specified concentration levels (Tsankov et al. 2015a; Tsankov et al. 2015b).

### 2.2. Assessing characteristics of differentiated hiPSC with BioQC

We applied the *BioQC* software developed previously to characterize the identity of the differentiated samples across all treated compound concentrations (including vehicle controls) on day 7 (Zhang et al. 2017). We used raw data of gene expression derived from molecular phenotyping and compared these profiles with tissue-preferential gene signatures derived from organ, tissue, and cell-type-specific gene expression data compiled from public compendia (Ljosa et al. 2013; Young et al. 2008). The BioQC performs Wilcoxon-Mann-Whitney tests comparing expression of genes in a set, e.g., genes preferentially expressed in one tissue, versus genes that are not in the set. The enrichment scores (log-10 transformed P-values) reported by BioQC are used to assess the similarity between the expression profile of interest and cell-type- and tissue-specific expression profiles.

### 2.3. Analysis and modeling of the *TeraTox* data

We performed differential gene expression (DGE) analysis comparing compound-treated samples with DMSO controls using the generalized linear model implemented in the *edgeR* package in R/Bioconductor (Robinson et al. 2009). To generate features for machine-learning models, we transformed the *P*-values associated with the coefficients of compound treatment to z-scores by the inverse of the quantile function of Gaussian distribution, multiplied by the sign of log2 fold-change (logFC). The vectors of *z*-scores of all genes (N=1,215) were used as raw features for machine-learning models, based on which further feature selection and engineering work was performed. We also tested the possibility of using the effect size, logFC, as a feature.

Besides the raw feature set of *z*-scores of all genes, we used three knowledge- and data-driven approaches to engineer the features in order to improve the performance of the machine-learning algorithms. First, we confined ourselves to the subset of germ-layer genes, because our and other’s work confirmed that their expression is specific to germ layers of embryogenesis, and their expression is modulated by teratogenic compounds (Suppl. Tab. S4) (Bock et al. 2011; Jaklin et al. 2020; Tsankov et al. 2015a; Tsankov et al. 2015b). Second, we used the germ-layer associations reported by Tsankov *et al.* to derive a reduced feature set defined by five germ-layer classes, including both germ layers (ectoderm, endoderm, mesoderm, mesendoderm) and pluripotency, by taking the median *z-*scores of germ-layer genes associated with each germ-layer class (Tsankov et al. 2015a). Finally, we used factor analysis, a dimension-reduction approach that derives latent variables from the correlation structure of observed variables, to identify latent biological, germ-layer factors (germ-layer factors for short), which reflect linear combinations of transcription factors, epigenetics, and other gene regulatory mechanisms that control embryogenesis.

We predicted teratogenicity potential in two ways. One way was to treat teratogenicity as a binary variable and to perform binary classification. The other way was to convert concentration-response teratogenicity into numeric metrics and to construct regression models. For the latter case, we defined a compound-specific Teratogenicity Score (TS hereafter). For non-teratogens, the TS was defined as 0 independent of the tested concentration. For teratogens, the TS was defined as the 0-1-bounded cosine similarity between the differential expression profile induced by a given concentration of a compound and the differential expression profile induced by the highest non-cytotoxic concentration of the same compound. The non-cytotoxic concentration was determined as the highest tested concentration associated with an average viability equal or larger than 80%.

The models were trained and validated using the Leave-One-Out (LOO) scheme. The full panel of compounds was assessed successively, leaving out one compound at a time and then used to build machine-learning models. We then compared teratogenicity scores predicted by the models with the observation of each left-out compound using the Spearman correlation coefficient. As an alternative to LOO, we also assessed repeated 80%/20% splitting of data into training sets and test sets.

In short, we considered two types of features (*z-*scores and logFC), four sets of factors (all genes / germ-layer genes / median *z*-scores or logFC of germ-layer classes defined by Tsankov *et al.* / median *z-*scores or logFC of germ-layer factors defined by factor analysis), two methods (linear regression with elastic net regularization and random forest, implemented in the *caret* package, version 6.0-88), two types of target variables (binary classification and regression), and two training/testing schemes (LOO and 80%/20% splitting). We tested all combinations exhaustively to build machine-learning models for teratogenicity scores and identified the best-performing models.

Besides predicting teratogenicity scores, we also comprehensively probed all options to build regression models for cytotoxicity (100%-viability), which was measured as part of the *TeraTox* assay. The same set of model architectures was tested, however, the combinations giving best performing models differed from that for teratogenicity scores (further discussed in results). All data analysis was performed with R (version 4.0.1) or Python (version 3.8.1) unless otherwise specified.

### 2.4. Test chemicals for validation

In total, we tested 28 positive and 17 negative reference substances in six-point concentrations in the mEST (see Suppl. Material and Methods, Suppl. Tab. S1) and the human *TeraTox* assay (Tab. 1). This compound panel consisted of both commercial and developmental pharmaceuticals with known teratogenicity potential (*i.e.* positive or negative) available from either human data, as reported in FDA drug labels, or from *in vivo* EFD studies in rats and/ or rabbits (ICH 2020). Some compounds without existing human or *in vivo* animal data were classified as teratogens based on a known teratogenic hazard associated with their mode of action (Belair et al. 2020; Chen et al. 2002; Cusack et al. 2017; Evans 2007; Kameoka et al. 2013; Lipinski et al. 2008; Sakata and Chen 2011; Wang et al. 2013; Worley et al. 2018). Compounds that did not result in increased incidences of birth defects in an adequate prospective cohort study accepted by health authorities were considered as non-teratogenic in humans, at least at the therapeutically-relevant exposure levels (Adams et al. 1969; Daniel et al. 2019; Dashe and Gilstrap 1997; Etwel et al. 2014; Muanda et al. 2017; Rumbold et al. 2015) (Suppl. Tab. S2).

The commercial compounds were obtained from Merck, Germany. The 12 investigational small molecule drug candidates RO-1 to RO-12 were provided by F. Hoffmann – La Roche, Switzerland (compound structures are not disclosed due to confidentiality and intellectual property issues). No human pregnancy data was available for the investigational drug candidates, but *in vivo* data were available from EFD studies in rats, and/or in rabbits (Suppl. Tab. S3). RO-1, RO-3, RO-8, RO-9 and RO-10 were teratogenic in EFD studies; RO-2, RO-4, RO-5, RO-6, RO-7, RO-11, RO-12 did not induce teratogenicity (Sergejew 2015).

All compounds were serially diluted in DMSO (0.25%) from a stock solution to six test concentrations and tested at appropriate non-cytotoxic concentration ranges in the *TeraTox* and *mEST* assays. We used the following metrics to compare the performance of the *TeraTox* and *mEST* assays. Sensitivity was calculated as the proportion of correctly predicted teratogens. Assay specificity was calculated as the proportion of correctly predicted non-teratogens. Overall accuracy was taken as the proportion of all correct predictions. F1 scores were calculated as the harmonic mean of precision and recall. True Positive, True Negative, False Positive, and False Negative are denoted with *TP*, *TN*, *FP*, and *FN*, respectively, and the performance metrics are defined in Supplementary Equations 1-5. To identify the threshold of TS that maximizes the performance (F1 score) of the *TeraTox Score*, we used a grid search between 0 and 1 with a step size of 0.01. The best threshold (TS=0.38) was chosen manually by inspecting the performance metrics.

To benchmark the performance of *TeraTox*, we applied two regulatory-accepted structure-based models to predict teratogenicity of commercially available compounds: the CAESAR model (version 2.1.8, Cassano et al., 2010) and the P&G model (version 1.1.2, Wu et al., 2013) implemented in the VEGA platform (version 1.3.10, Marzo et al., 2016). For the benchmark, we used 20 compounds (15 teratogens and 5 non-teratogens) that were not part of the training set of the CAESAR model.

### 2.5. Model explainability and interpretation

We used the Type I importance measure of features (mean decrease in accuracy) of random-forest models to compare the importance of germ-layer genes in the teratogenicity model and in the cytotoxicity model.

Pharmacology data of publicly available compounds were downloaded from ChEMBL (version 26). We only used human targets and affinities derived from high-quality dose-response data. Binary distances were used to cluster the compounds by their pharmacological profiles.

To construct a Bayesian network model of regulations between factors, we first discretized differential gene expression data of the first six germ-layer factors into three levels using the Hartemink’s pairwise mutual information method implemented in the *bnlearn* package (Scutari 2010). We generated 1,000 bootstrap replicates using Hill Climbing, a score-based learning algorithm, and the Bayesian Dirichlet equivalent (uniform) score (bde, with the imaginary sample size set to 10). Edges that persisted in more than 85% bootstrap samples were deemed as significant and reported.

The beta regression model used for sensitivity analysis was built with the *glmmTMB* package (Brooks et al. 2017). Scores outside the boundaries [0.01, 0.99] are set to the boundary values to allow beta regression. All ten factors and significant interaction terms identified in the Bayesian network were used as the model input and compounds were modelled as random effects to capture between-concentration correlations. For better interpretability, input variables were scaled to zero mean and standard deviation. Simulation was performed using the *ggeffects* package (Lüdecke 2018).

## 3. Results

### 3.1. Gene expression quantification by molecular phenotyping

We previously described that differential expression of a set of 87 genes preferentially expressed in different germ layers is in principle able to distinguish between teratogenic and non-teratogenic compounds (Jaklin et al. 2020). These germ-layer genes both determine and reflect embryonic development (Tsankov et al. 2015a). To validate our findings, we compiled a large set of well-documented teratogens and non-teratogens that are challenging to predict and/or known to cause false-positives in animal studies (Suppl. Tab. S2, S3). The compounds cover a broad spectrum of chemical classes and a wide range of effective concentrations.

We evaluated the performance of our human stem-cell model by testing the panel of compounds, adapting the experimental workflow developed previously (Fig. 1a and 1b). We identified the assay throughput as a major challenge due to the high number of samples for gene expression profiling (>1,000). It would be particularly cost- and labor-intensive to use the digital PCR technique established in our previous work to quantify gene expression (Jaklin et al. 2020). To address this challenge, we used molecular phenotyping as an alternative readout. Molecular phenotyping is based on amplicon-based targeted sequencing and is able to deliver quantitative expression data of 1,215 pre-defined genes, including both pathway reporter genes, *i.e.*, genes that are specifically modulated by pathway perturbations, as well as germ-layer genes that we reported in our previous study. In this way, we were able to characterize both general pathway activity modulations and germ layer-specific changes as potential features associated with teratogenicity (Drawnel et al. 2017; Zhang et al. 2015; Zhang et al. 2014).

**Figure 1:**
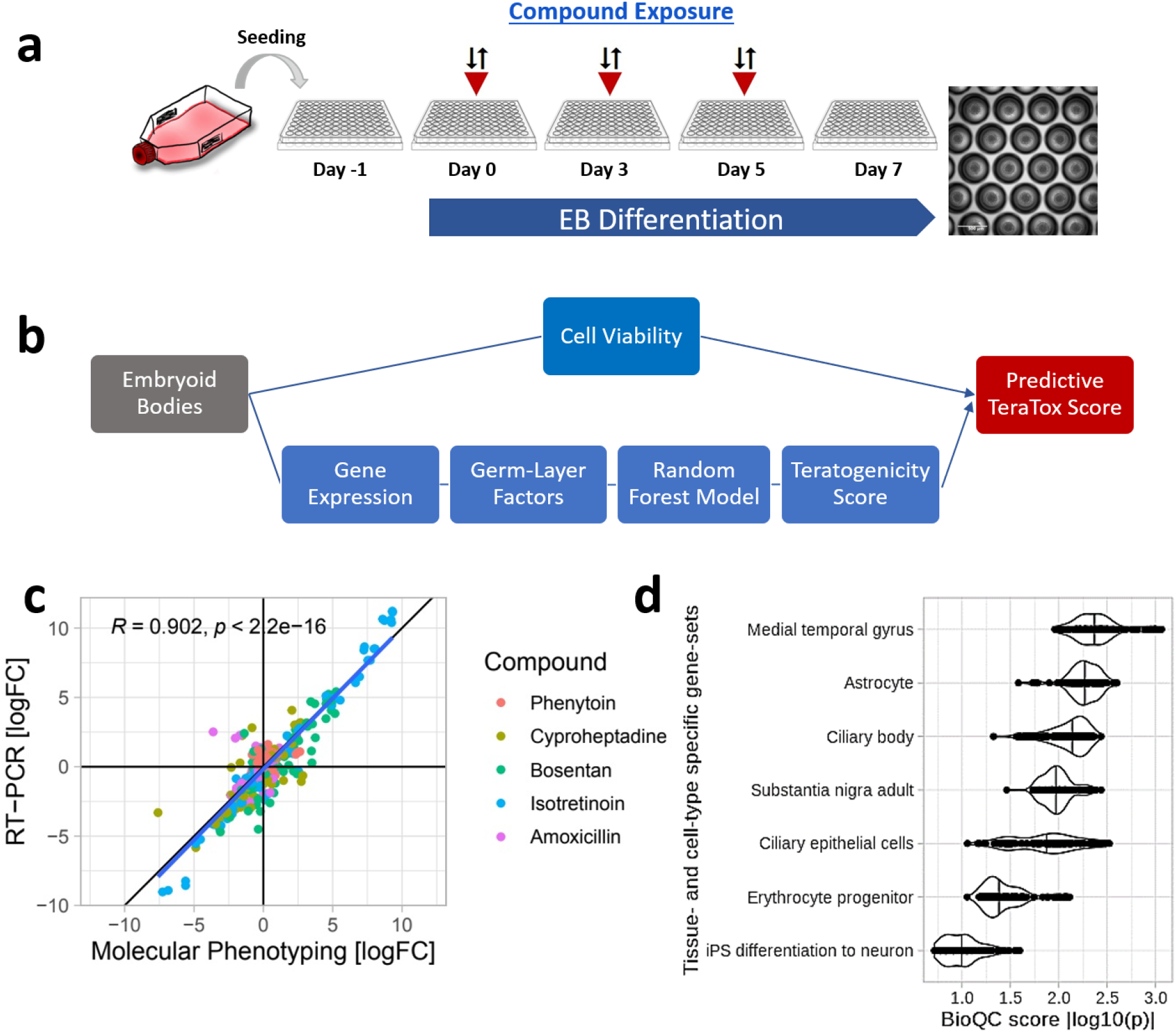
The human *TeraTox* assay: workflow and quality control. **(a)** hiPSC differentiate over 7 days and form EBs. Compounds are added on day 0, 3 and 5. Single wells of differentiated EBs are lysed for one sample. **(b)** *TeraTox* score is calculated based on cell viability, gene expression data, and machine-learning model. **(c)** DGE from molecular phenotyping correlated with data from RT-PCR assay represented in log2 fold change (logFC). Dots represent germ-layer genes. *R* = Spearman correlation coefficient between two sets of measurements in all compounds. **(d)** *BioQC* of raw gene expression data (DMSO controls) revealed the biological identity of hiPSC and showed significantly enriched cell-type signatures (median p<0.10). Each dot = one sample. Violins show distributions of BioQC scores (absolute log10 transformed *p-*values of the Wilcoxon-Mann-Whitney test) from each gene set, vertical lines indicate median values. The larger the BioQC score, the more enriched is the expression of the genes.

We performed extensive quality control of the data. Here we address the questions whether results of molecular phenotyping are comparable with those of qRT-PCR, and whether the hiPSC used show expected cell identity based on their gene expression profile. We compared the differential expression profiles of germ-layer genes obtained by qRT-PCR in previous studies with newly generated data of molecular phenotyping and observed highly consistent results (Pearson correlation coefficient R=0.9, p<2.2E16) (Fig. 1c). Molecular phenotyping requires far fewer cells and delivers much higher throughput than qRT-PCR, allowing a marked improvement in the productivity of the *TeraTox* assay. A unique advantage of quantifying pathway reporter genes along with germ-layer genes is the identification of cell-type-specific gene expression patterns. To this end, we applied *BioQC* analysis, a method that we developed to identify sample heterogeneity and tissue comparability using gene sets preferentially expressed in cells and tissues (Zhang et al. 2017). We observed that the expression profiles of the cells used in the *TeraTox* assay at day 7 resemble a mix of those gene signatures specific for astrocytes, epithelial cells, and iPSC derived neurons (Fig. 1d). This suggests that the hiPSC used for the assay have a preferred differentiation propensity into the neuroectodermal lineage, which is in agreement with previous time-series gene expression studies that demonstrated pronounced expression of ectodermal markers at day 7, followed by meso- and endodermal expression (Jaklin et al. 2020; Tsankov et al. 2015a).

### 3.2. Unsupervised learning from gene expression data with factor analysis

Before applying supervised learning techniques to differentiate teratogens from non-teratogens, we applied several unsupervised learning algorithms to explore the gene expression data, including principal component analysis (PCA) and factor analysis. PCA revealed experimental plate effects that we could successfully correct with linear regression models for differential gene expression (data not shown). Unexpectedly, factor analysis revealed both biological insights and suggested a technique for feature engineering to produce the best-performing model (see below, also a brief introduction to factor analysis is given in Supplementary Material and Methods).

We applied factor analysis to raw gene expression data and identified intriguing patterns. Since factor analysis is based on inter-gene correlations, we visualized the correlation matrix of germ-layer genes in Figure 2a (the full matrix is visualized in Supplementary Figure S2a). Genes that strongly correlate with each other form clusters, which correspond to latent factors.

**Figure 2:**
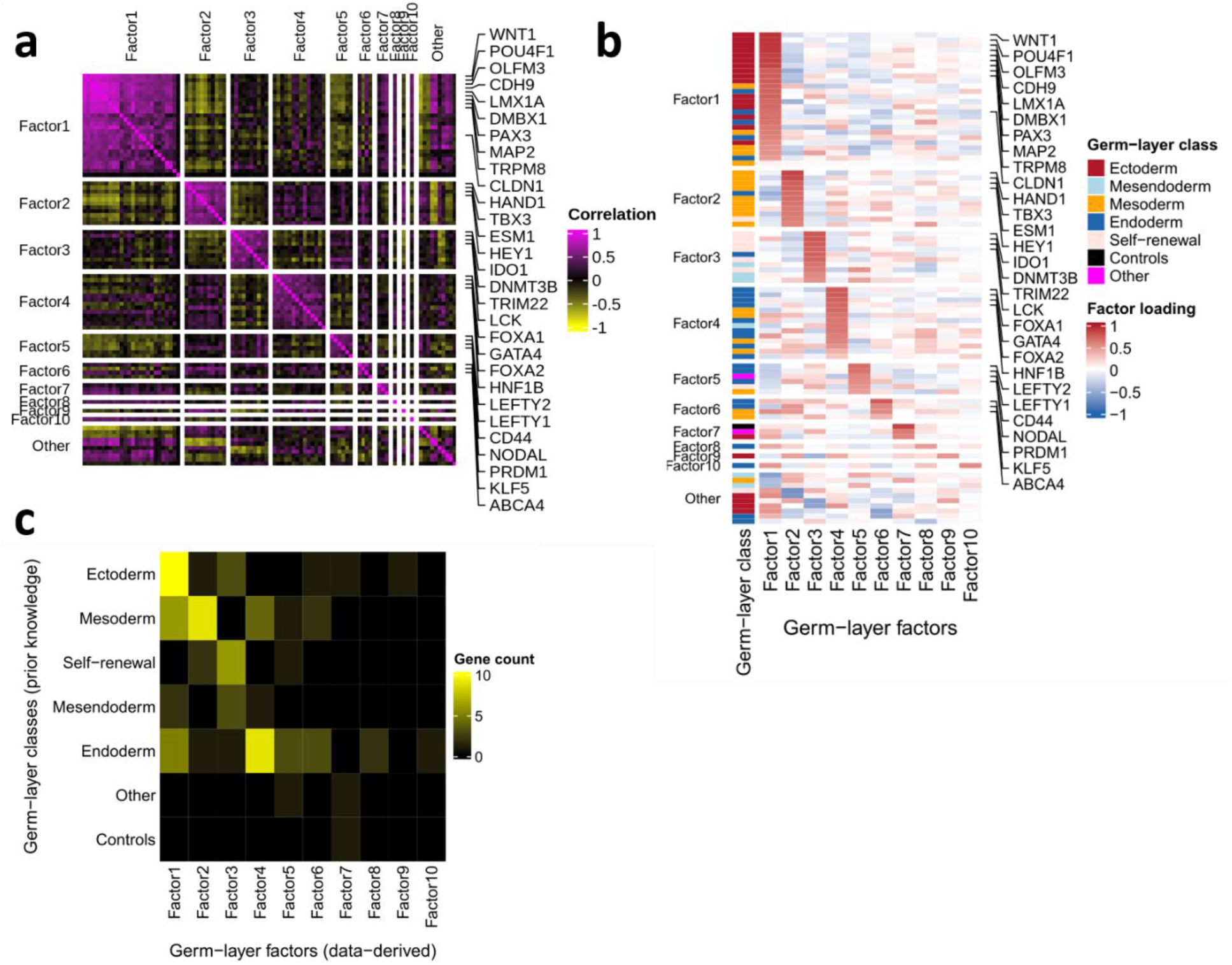
Identification of latent factors that are associated with germ layers. **(a)** Germ-layer genes show correlated expression and form clusters of co-expression. The heatmap represents pairwise Pearson correlation coefficients of germ-layer gene expression in all samples. Genes are split by latent germ-layer factors for representative genes (full matrix is shown in Supplementary Figure S2a). **(b)** Loadings of factor analysis (germ-layer genes in rows and linear combinations of latent germ-layer factors in columns). Loadings equal to or near +1 or −1 indicate that the factor positively or negatively influences the gene, while loadings near 0 means that the factor has little effect on the gene (full matrix in Suppl. Fig. S2c). **(c)** Germ-layer factors are not equivalent to, but significantly associated with, germ-layer classes. The heatmap visualizes the number of genes shared by each pair of germ-layer classes (in rows) and germ-layer factors (in columns).

Despite that factor analysis is a correlation-based statistical method in which we injected no prior knowledge, it revealed biologically meaningful patterns. Using the maximum likelihood method, we decomposed the covariance matrix of gene expression into factors. The heatmap in Figure 2b shows loadings, *i.e.* how strong factors influence the expression of germ-layer genes, of the first ten factors that collectively explain more than 70% of the covariance (Suppl. Fig. S2b and S2c). Left to the heatmap we use colors to indicate germ-layer classes that were distilled from biological knowledge. We found that the first six factors (ranked by explained covariance of the data) are significantly enriched with signatures of individual germ layers or signatures of stem-cell self-renewal (Fig. 2c, p<0.01, Fisher’s exact test). This significant enrichment is intriguing, because while it is established that germ-layer genes are highly expressed at different stages of embryogenesis, factor analysis reveals for the first time that their expressions are strongly correlated in 3D embryoid bodies formed by hiPSC, with or without compound treatment. Given that the cells used in *TeraTox* are cultured up to day 7, it is unlikely that the correlations are caused by temporal changes of embryogenesis. Instead, factor analysis suggests that besides being correlated across time in development, expression of germ-layer genes is also correlated across treatment conditions in 7-day spontaneously differentiated EBs.

Detailed analysis of the results from the factor analysis revealed more insights. The strongest correlation of the germ-layer genes was observed among genes in Factor 1, many of which are markers of the ectodermal layer, *e.g.*, *WNT1*, *POU4F1*, *OLFM3*, *CDH9*, *LMX1A*, *DMBX1*, *PAX3*, *MAP2*, and *TRPM8* (Fig. 2a). While BioQC analysis revealed that ectodermal genes are highly expressed at the endpoint on day 7, factor analysis further indicated that their expression is strongly correlated across conditions, too, which is neither sufficient nor necessary for their high expression. Factors 2-6 mainly consist of genes representing the mesodermal layer (Factor 2), stem-cell self-renewal (Factor 3), and the endoderm layer (Factor 4-6), respectively. The remaining factors (Factor 7-10) are of smaller sizes and more heterogeneous (Fig. 2b). Genes associated with each factor are associated mainly, but not exclusively, with other genes of the same germ-layer class.

In summary, factor analysis revealed that germ-layer genes form co-regulated gene modules in *TeraTox* that are significantly enriched by germ-layer- or stem-cell-specific markers.

### 3.3. Training and testing of a predictive model for the *TeraTox* assay

To build a quantitative predictive model of concentration-dependent teratogenicity potential with gene expression, we explored all combinations of the following options (Fig. 3a):

1. *Feature type*: We tested both log2 fold change (logFC), the point-estimate of the effect size, and z-scores transformed from the sign of logFC and *p*-value reported by the *edgeR* model, which considers both effect size and variance of differential gene expression.
2. *Feature engineering*: We used all detectable pathway reporter genes (N=1,215), detectable germ-layer genes (N=87), germ-layer classes defined by Tsankov *et al*. (N=7), and germ-layer factors derived from factor analysis (N=10). For both germ-layer classes and factors, we used the median value of the genes belonging to each group as the engineered feature.
3. *Model construction*: We used and benchmarked two methods of different nature, Elastic Net (linear regression with regularization) and Random Forest (ensemble decision trees), to construct machine-learning models. These methods were chosen based on the size of the dataset and the relatively good explainability of both methods (Badillo et al. 2020).
4. *Target variable*: We used both binary classification (teratogen or non-teratogen) and regression (the teratogenicity score, defined below and further detailed in the Material and Methods section) for teratogenicity and regression alone for cytotoxicity.
5. *Data splitting*: we tried both repeated splitting of 80% training and 20% test set, and the leave-one-out (LOO) scheme. For data splitting, we used 80% of compounds (stratified sampling from non-teratogens and teratogens) as the training set to train a model, which was used to predict the teratogenicity scores using the remaining 20% compounds as the test set. For LOO, the model was trained by assessing the panel of compounds minus one, which predicted the teratogenicity scores for the left-out compound. The procedure was repeated until all compounds had been left out. The performance of both models was assessed by F1 scores in case of binary classification models, and Spearman correlation coefficients of teratogenicity scores for teratogens in case of regression models. The best model parameters were searched by 10-fold cross-validations of the training set.

**Figure 3:**
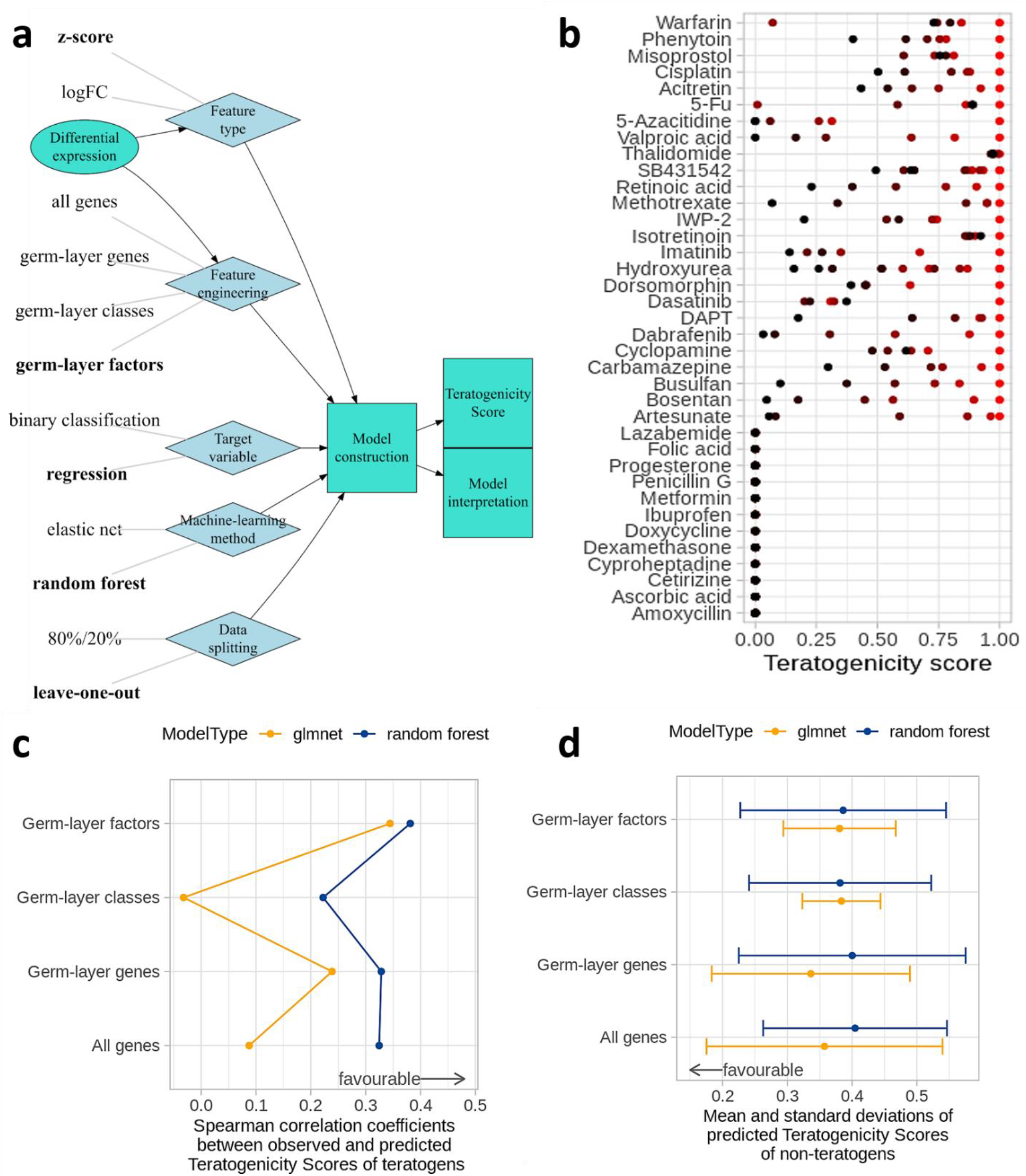
Construction of machine-learning models predicting concentration-dependent teratogenicity potentials based on differential gene expression as input. **(a)** Overview of investigated model architectures. **(b)** Definition of teratogenicity scores (TS) from highest (red) to lowest (black) concentrations. TS of teratogens = 1 for the highest non-cytotoxic concentration, TS for other concentrations = cosine similarity of differential gene expression profiles between each concentration and the highest non-cytotoxic concentration. TS of non-teratogens = 0, independent of the concentration level (from *Lazabemide* to *Amoxicillin*). Negative TS = 0. **(c)** Spearman correlation coefficients between observed teratogenicity scores, calculated on a per-compound basis, and predicted teratogenicity scores, which are derived from models trained by leave-one-out testing. **(d)** Mean (dots) and standard deviations (error bars) of teratogenicity scores of non-teratogens. Median teratogenicity score of each compound is derived from six concentrations. The average scores of non-teratogens are lower than those of teratogens, but not strictly zero, because they are predicted values by leave-one-out testing with our machine learning model instead of assigned values as in (b).

The teratogenicity scores of teratogens are defined between 0 and 1, and those of non-teratogens are fixed as 0 at all concentrations (Fig. 3b). By defining teratogenicity scores, we effectively transformed the binary classification problem into a regression problem. We refer readers interested in the motivation of developing the Teratogenicity Score and in the mathematical details to the section on Teratogenicity Score in Supplementary Material and Methods.

We observed the following patterns as we tried all options of model building:

1. The *feature type* has minimal impact on the performance, though models trained with *z-*scores perform better on the test set than models trained with logFC (data not shown).
2. The combination of *feature engineering* and *machine-learning model* is important and the best combination depends on the prediction task (Fig. 3c and 3d). For teratogenicity prediction, the combination of germ-layer factors and random-forest regression worked best.
3. With regard to the *target variable*, the performance of the regression-based teratogenicity-score prediction model is slightly better than binary classification (data not shown).
4. Performance is comparable between two modes of *data splitting* (data not shown). However, the LOO training-testing scheme is preferable because it allows us to set up a single threshold of teratogenicity score, which can be applied to all compounds, and is not conditioned by whether or not a compound is included in the training set or in the test set as in the case of 80%/20% data splitting.

Based on these observations, we decided to use germ-layer factors as features, random-forest regression as the machine-learning model, and teratogenicity score as the target variable to build the predictive model for teratogenicity with gene expression data.

### 3.4. Performance of the *TeraTox* assay and benchmarking with other models

Based on the best-performing machine-learning model, we defined the following predictive model for teratogenicity. First, we considered the maximal non-cytotoxic threshold concentration (NCC_max_) for cell viability of at least 80%, measured by the CellTiter Glo assay. Next, we defined the minimal teratogenic concentration (TC_min_) as the concentration at which the threshold of the teratogenicity score was met (TS=0.38, defined by grid search, Fig. 4a). If no NCC_max_ or TC_min_ could be determined because values did not exceed these thresholds, the maximal tested concentrations were used for NCC_max_ and TC_min_. The predictive score, which we named *TeraTox Score,* is defined by the logarithmic ratio between threshold concentrations at 20% viability impairment (NCC_max_) and teratogenic concentrations (TC_min_). Negative *TeraTox* scores classify the compounds as negative whereas positive scores classify compounds as positive (Fig. 4b).

**Figure 4:**
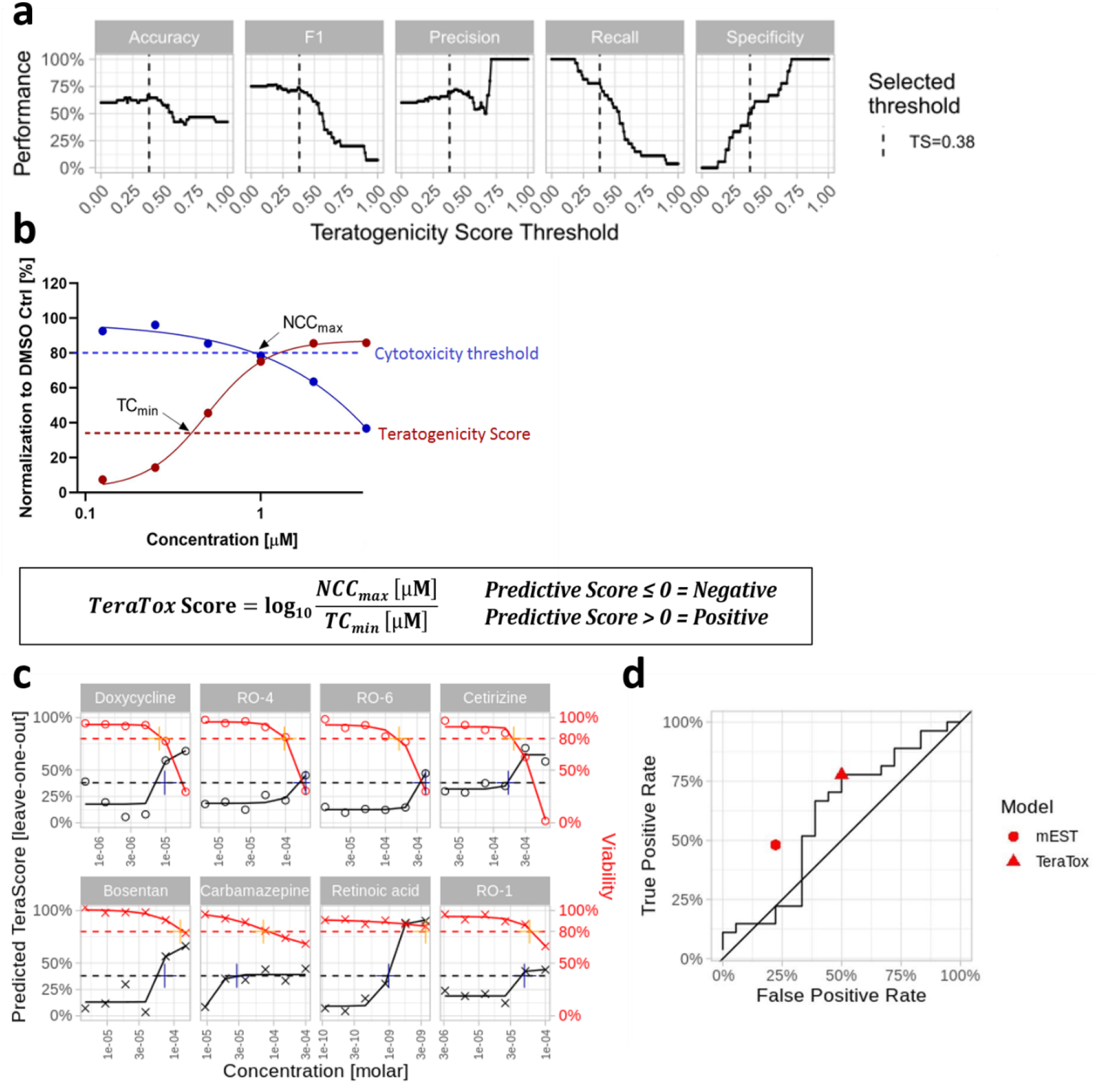
Prediction of teratogenicity with the human *TeraTox* assay. **(a)** Results of a grid search to select the optimal threshold of the teratogenicity score using the best identified model architecture. The best threshold (TS=0.38) was chosen based on performance metrics defined in Suppl. Equations (1)-(5). **(b)** Visual definition of the *TeraTox* score based on minimal teratogenic concentration (TC_min_) and maximal non-cytotoxic concentration (NCC_max_). Compounds with *TeraTox* scores ≤0 are classified as negative and compounds with scores >0 are classified as positive. **(c)** Examples of concentration-response curves reported by the *TeraTox* assay of four selected non-teratogens (top panels) and four selected teratogens (bottom panels). The “+” indicates predicted teratogenicity score (n=2) and measured cytotoxicity (n=3). **(d)** Receiver operating characteristics (ROC) curve of leaving-one-out tests based on 45 reference compounds.

We plotted the concentration-response curves of measured cytotoxicity and predicted teratogenicity scores induced by each compound (Fig. 4c, see Suppl. Fig. S4 for all compounds). In general, teratogenicity levels increased while cell viability decreased with rising concentrations. Correctly predicted negative compounds were unlikely to induce teratogenicity within non-cytotoxic concentrations, which means the calculated *TeraTox* score was negative or zero (e.g., Doxycycline, RO-4, RO-6). Positive compounds (e.g., Bosentan, Carbamazepine, Retinoic Acid, RO-1) or false positive predicted compounds (e.g., Cetirizine) were more likely to induce teratogenicity under non-cytotoxic concentrations, as indicated by positive *TeraTox* scores (Fig. 4c).

We compared the *TeraTox* prediction scores with classifications from FDA or *in vivo* EFD studies for 45 reference compounds (Suppl. Tab. S5). Classification with *TeraTox* Scores achieved an overall accuracy of 69% and outperformed mEST (58%). The two assays show different sensitivity and specificity profiles: While mEST is more specific (specificity 76%), *TeraTox* is more sensitive (sensitivity/recall 79%). Among 17 negative reference compounds, 8 were classified as false positives (FP) by *TeraTox*, and only 4 by the mEST. Whereas from 28 positive reference compounds, 22 were predicted as true positives (TP) by *TeraTox* and only 13 by the mEST (Tab. 2, Fig. 4d). It is noteworthy that among the 26 compounds misclassified in total, the following seven compounds are wrongly predicted by both assays: cyproheptadine, RO-11, 5-FU, methotrexate, misoprostol, RO-8, and warfarin. Given the distinct sensitivity and specificity profiles of the two assays, we asked whether we can achieve even better prediction results by using the two tests in sequence. Therefore, if we first run the mEST on the full panel, the substances with negative mEST results would then be retested by *TeraTox* to benefit from the high specificity of mEST and the high sensitivity of *TeraTox*. Indeed, we found that overall accuracy of the combined prediction increased to 78%, better than either *TeraTox* or *mEST* alone. This suggests that it may be possible to achieve better prediction results by combining the existing mEST assay with the novel *TeraTox* assay.

Furthermore, we compared 18 pharmaceutical compounds that were both tested in *TeraTox* and *DevTox*^*qp*^ by Stemina, and observed the identical accuracy (78%), whereas balanced accuracy was 73% for *TeraTox* and 87% for *DevTox*^*qp*^ assay. The *DevTox*^*qp*^ assay delivered a higher specificity (100%) compared to *TeraTox* (67%), whereas *TeraTox* was more sensitive (80%) than *DevTox*^*qp*^ (73%) (Supplementary Tab. S6 & S7).

We also compared *TeraTox* with in *silico* predictions of developmental and reproductive toxicity using two widely used QSAR models: CAESAR and P&G, both implemented in the VEGA software (Suppl. Tab. S8). Among 20 compounds that were not used as training sets for CAESAR, we found that *TeraTox* performed better than both the CAESAR model (85% versus 75%) and the P&G model (accuracy of 52%).

In short, *TeraTox* delivers comparable or more favorable performance with alternative assays, and combining *TeraTox* with other assays can further increase the prediction accuracy for drug-induced teratogenicity.

### 3.5. Leveraging *TeraTox* data and model to gain biological insights

A model’s explainability is crucial for its understanding to allow for inspection and further improvement (Barredo Arrieta et al. 2020). We performed additional in-depth analysis of the cytotoxicity and gene expression data and collected additional data orthogonal to *TeraTox*, thereby implementing four independent approaches to interpret and explain how the machine-learning model works and to explore what differs teratogens from non-teratogens.

First, we followed up on previous work and asked the question whether compound-induced cytotoxicity quantified by the phenotypic assay can be predicted by gene expression data as well, and whether teratogenicity scores are confounded by general cytotoxicity (Waldmann et al. 2016; Waldmann et al. 2014). For this purpose, we followed the same scheme as described in Figure 3a using cytotoxicity instead of teratogenicity scores as the target variable. Interestingly, a comprehensive search showed that using all pathway reporter genes and the *elastic net* model, instead of using germ-layer factors and random forest as in the case of teratogenicity prediction, give the best result (Fig. 5a, contrasted with Fig. 3c and 3d).

**Figure 5:**
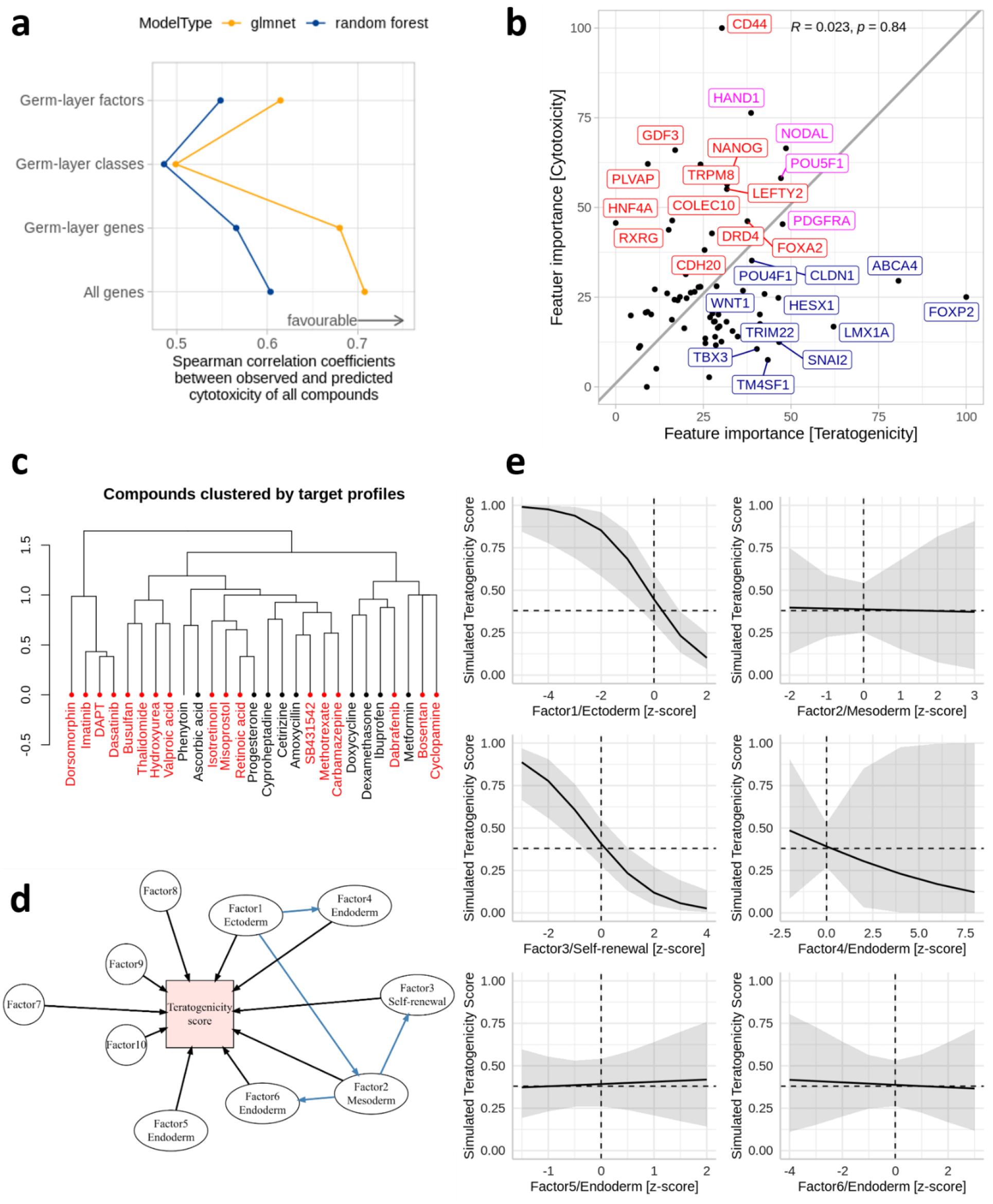
Biological interpretation of the model. **(a)** Prediction of cytotoxicity based on DGE (same workflow as Fig. 3a) using all pathway reporter genes as features and an elastic net machine-learning model. Dots: Spearman correlations between observed and predicted cytotoxicity using LOO testing. **(b)** Difference in feature importance of germ-layer genes for prediction of cytotoxicity (red) or teratogenicity (blue) or both (magenta). **(c)** Clustering analysis: pharmaceutical target profiles of compounds alone are not sufficient to determine teratogenicity (based on annotated compound target profile, *i.e.* quantitative affinity to protein targets from ChEMBL database). Teratogens = red, non-teratogens = black. **(d)** Structure of the Directed Acyclic Graph (DAG) to model relationships between teratogenicity score and germ-layer factors with a generalized linear model. Input variables: germ-layer factors and significant interactions between germ-layers identified by Bayesian networks (blue edges). Model fitting: Suppl. Fig. S6. **(e)** Generalized linear model with beta-regression for sensitivity analysis, to test model behavior with tuned input variables. Each panel shows the result of one tuning parameter, e.g. ectoderm germ-layer factor (top-left) while keeping all other parameters fixed. Black lines = average prediction, gray areas = 95% confidence intervals of prediction. Input variables are scaled to 0 mean and standard deviation.

Given that the combination of germ-layer genes and random forest gives reasonable performance in both cases, and that random forest allows inquiry of feature importance by accuracy, we compared the feature importance of germ-layer genes in predicting both target variables (Fig. 5b). The prediction of cytotoxicity and teratogenicity by molecular phenotyping relies on expression changes of distinct genes. The distinction shows that teratogenicity of a compound is not a determinant for cytotoxicity whereas a compound that shows cytotoxicity at a specific concentration can still be teratogenic at lower concentrations and that pathways for cytotoxicity and teratogenicity may be independently regulated. This concurs with several previous findings (Krug et al. 2013; Rempel et al. 2015; Shinde et al. 2017).

The second approach addressed the question whether a compound’s pharmacology, namely its target profile (protein targets and binding affinities), suffices to predict its teratogenicity potential. If so, one may hope to predict teratogenicity potential based on target profiles and/or even based on the chemical structure alone. While some teratogens indeed have similar target profiles, we observe close clustering of teratogens and non-teratogens that have similar target profiles as well (Fig. 5c, Suppl. Fig. S5a). The potential of teratogenicity, therefore, may be associated with off-target effects or effects through targets that are not captured in ChEMBL, especially at the relatively high concentrations approaching cytotoxicity levels that we tested. Corroborating this, we found almost no correspondence between clustering of average differential gene expression across concentration per compound and that of pharmacological profile (Suppl. Fig. S5b). Therefore, we conclude that while knowing the target- and off-target profile of a compound is essential for de-risking its safety liabilities including teratogenicity, pharmacology data alone cannot currently predict a compound’s teratogenicity potential. This conclusion concurs with the superior performance of *TeraTox* over two QSAR models, which consider the chemical structure alone. Human iPSC based *in-vitro* assays, such as *TeraTox* and other advanced cellular models, are therefore indispensable for assessing the potential for human teratogenicity.

The third approach was to use a simpler generalized linear regression model for sensitivity analysis, which would allow us to analyze how the model responds to changes of the input. Given that random forest is an ensemble method and the contribution of each germ-layer factor can be therefore difficult to interpret, we built an alternative model using beta linear regression. To identify interaction terms in the linear regression, we made the assumption that germ-layer factors regulate each other by forming a directed acyclic graph (DAG). Under this assumption, we built a Bayesian network using the differential expression data of germ-layer factors (Fig. 5d). The network reveals potential influences on both mesoderm and endoderm by the ectoderm, influences on endoderm by mesoderm, and influences on stem-cell renewal by endoderm.

The Bayesian network topology prompted us to build a beta regression model including all germ-layer factors and interactions identified in the Bayesian network (Fig. 5e, Suppl. Fig. S6). The model provides both interpretable coefficients of the model and a tool for sensitivity analysis, because we can quantify prediction uncertainty much easier with a linear model than the random forest model, by paying the price of assuming linear regulation relationship. For the sensitivity analysis, we kept all other parameters fixed and tuned one input parameter at a time to simulate its impact on predicted teratogenicity scores. We observed that the model is likely sensitive to impairment of either ectoderm layer or stem-cell self-renewal, while being relatively robust to changes to either mesoderm or endoderm (Fig. 5e). The results of sensitivity analysis further underlined the prominent ectodermal nature of the model at the endpoint on day 7.

Last but not least, we applied gene-set enrichment analysis to each compound with *BioQC* and compared median gene-set enrichment results over concentrations of each compound between teratogens and non-teratogens. We identified multiple gene-sets that are potentially differentially regulated by teratogens and non-teratogens (p<0.10, Suppl. Figure S7). Interestingly, target genes of several transcription factors that are involved in organ development and cell differentiation, for example POU4F1, GATA1, and NODAL, were impacted by teratogens differently from non-teratogens. Besides, teratogens also regulate genes involved in biological processes that are not restricted to embryogenesis, such as cleavage of cell adhesion proteins as well as lipid metabolism genes induced by SRBEF/SREBP. While teratogens do not form a homogeneous group and have distinct pharmacological profiles (Fig. 5c, Suppl. Fig. S5a), these results suggest that transcriptional regulation by teratogens can manifest in changes of several biological processes, potentially mediated by key transcriptional factors. Gene-set enrichment analysis based on *TeraTox* data also revealed molecular insights of teratogenic effects at yet another level of gene expression regulation (Suppl. Fig. S7).

In summary, we explain how the *TeraTox* model operates by complementing the machine-learning model with feature importance analysis, biological and pharmacological interpretation, sensitivity analysis, and gene-set enrichment analysis.

## 4. Discussions

This study characterizes the optimization of *TeraTox* and its application in the context of drug development to detect teratogenicity. *TeraTox* extends and standardizes the previously published embryoid-body models and fully leverages the predictive potential of these models by adding a toxicological prediction model (Krug et al. 2013; Shinde et al. 2016). It exploits an explainable machine-learning approach to predict teratogenicity potential induced by drug-like molecules.

Only a few pharmaceutical compounds have been recognized as human teratogens based on high-quality data. Therefore, we decided to limit the training data of the *TeraTox* model to a compound set with well-described evidence without losing rigor in the teratogenicity classification (N=45, Table 1). While alternative compound collections are available, for instance the *ToxCast* data set applied to the *DevTox*^*qp*^ assay from Stemina (Zurlinden *et al.* 2020), our compilation was specifically related to pharmaceuticals.

**Table 1:** Reference compounds used for assay validation, with human teratogenicity classification and test concentration according to non-cytotoxic concentrations tested in the human *TeraTox* assay. (dilution ratios in brackets covering 6 concentrations). Teratogenicity classification was based on FDA classification (Suppl. Tab. S2) or *in vivo* EFD data (indicated with asterisks*, Suppl. Tab. S3).

| **Reference Compound** | **Teratogenicity Classification** | **Test Concentrations (human model) [μM]** |
| --- | --- | --- |
| **Acitretin** | **Positive** | 0.08 – 2.5 (1:2) |
| **Amoxicillin** | Negative | 6.25 – 200 (1:2) |
| **Artesunate** | **Positive** | 0.13 – 8 (1:2) |
| **Ascorbic Acid** | Negative | 28 – 900 (1:2) |
| **Bosentan** | **Positive** | 4.7 – 150 (1:2) |
| **Busulfan** | **Positive** | 0.06 – 4 (1:2) |
| **Carbamazepine** | **Positive** | 4.7 – 300 (1:2) |
| **Cetirizine** | Negative | 9.3 – 600 (1:2) |
| **Cyclopamine** | **Positive** | 0.3 – 20 (1:2) |
| **Cyproheptadine** | Negative | 0.47 – 30 (1:2) |
| **Dabrafenib** | **Positive** | 0.03 – 2 (1:2) |
| **DAPT** | **Positive** | 0.05 – 3 (1:2) |
| **Dasatinib** | **Positive** | 0.3 – 20 (1:2) |
| **Dexamethasone** | **Positive** | 4.7 – 300 (1:2) |
| **Dorsomorphin** | **Positive** | 0.2 – 14 (1:2) |
| **Doxycycline** | Negative | 0.3 – 20 (1:2) |
| **5-Fluorouracil** | **Positive** | 0.004 – 0.25 (1:2) |
| **Hydroxyurea** | **Positive** | 3.12 – 200 (1:2) |
| **Ibuprofen** | Negative | 1.9 – 1400 (1:3) |
| **Isotretinoin** | **Positive** | 4.7 – 300 (1:2) |
| **Imatinib** | **Positive** | 1.6 – 100 (1:2) |
| **IWP-2** | **Positive** | 0.0015 – 0.1 (1:2) |
| **Lazabemide** | Negative | 1.6 – 100 (1:2) |
| **Metformin** | Negative | 7.8 – 500 (1:2) |
| **Methotrexate** | **Positive** | 0.0025 – 40 (1:5) |
| **Misoprostol** | **Positive** | 0.02 – 1.3 (1:2) |
| **Penicillin G** | Negative | 9.3 – 600 (1:2) |
| **Progesterone** | Negative | 0.63 – 40 (1:2) |
| **Retinoic Acid** | **Positive** | 0.0005 – 0.035 (1:2) |
| **RO-1\*** | **Positive** | 1.6 – 100 (1:2) |
| **RO-2\*** | Negative | 7.8 – 500 (1:2) |
| **RO-3\*** | **Positive** | 4.7 – 300 (1:2) |
| **RO-4\*** | Negative | 3.1 – 200 (1:2) |
| **RO-5\*** | Negative | 0.8 – 50 (1:2) |
| **RO-6\*** | Negative | 6.25 – 400 (1:2) |
| **RO-7\*** | Negative | 9.3 – 600 (1:2) |
| **RO-8\*** | **Positive** | 1:25 – 80 (1:2) |
| **RO-9\*** | **Positive** | 0.08 – 5 (1:2) |
| **RO-10\*** | **Positive** | 0.23 – 15 (1:2) |
| **RO-11\*** | Negative | 0.6) – 40 (1:2) |
| **RO-12\*** | Negative | 1.6 – 100 (1:2) |
| **SB431542** | **Positive** | 0.31 – 20 (1:2) |
| **(±) Thalidomide** | **Positive** | 0.0007 – 0.5 (1:3) |
| **Valproic Acid** | **Positive** | 15.6 – 1000 (1:2) |
| **Warfarin** | **Positive** | 0.9– 60 (1:2) |

**Table 2:** Overview of assay performance for mEST and human *TeraTox*assay. Values were calculated based on 45 compounds (according to supplementary equations 1-5. TP= true positive, TN=true negative, FP=false positive, FN=false negative).

| ***Model*** | ***TP*** | ***TN*** | ***FP*** | ***FN*** | ***Accuracy*** | ***Precision*** | ***Recall*** | ***Specificity*** | ***F<sub>1</sub>*** |
| --- | --- | --- | --- | --- | --- | --- | --- | --- | --- |
| ***TeraTox*** | 22 | 9 | 8 | 6 | 69% | 73% | 79% | 53% | 76% |
| ***mEST*** | 13 | 13 | 4 | 15 | 58% | 76% | 46% | 76% | 57% |
| ***mEST +***<br>***TeraTox*** | 22 | 13 | 4 | 6 | 78% | 85% | 79% | 76% | 82% |

Despite the limited number of reference compounds available, we report that *TeraTox* outperforms two QSAR models and performs comparably with state-of-the-art *in-vitro* models. *TeraTox* and *DevTox*^*qp*^ showed the same accuracy to predict 18 pharmaceutical compounds: the *DevTox*^*qp*^ assay showed a higher specificity and *TeraTox* a higher sensitivity. Combining *TeraTox* and *DevTox*^*qp*^ led to a higher prediction accuracy, suggesting a synergistic effect of using complementary assays for teratogenicity prediction.

Similarly, we observed comparable performance of *TeraTox* and mEST, with *TeraTox* being more sensitive and mEST more specific. Again, combining both assays led even to an increased accuracy, which would improve overall predictivity.

Due to our long experience with mEST assay, we could also offer a first-hand comparison between *TeraTox* and mEST with regard to manual work, cost, and throughput (Suppl. Tab. S9). We conclude that the overall effort and cost entailed by *TeraTox* is comparable to that of the mEST. However, *TeraTox* offers a much higher throughput thanks to automation and miniaturization, and it generates quantitative gene expression data that can be used to compare new chemical series with existing ones and to further refine the model.

The prediction model presented here is geared towards hazard identification, similar to animal studies, where maximum tolerated doses are to be used for studies testing DART. In the context of an overall risk assessment, one immediate step would be to consider compound potency and to evaluate whether the *in vitro* toxic concentrations would be relevant to human exposure situations. To exemplify this, we considered the false positives identified here (Suppl. Tab. S2 & S5).

Two of them have a minimal transcriptome-altering concentration that is > 2 orders of magnitude higher than the human max plasma concentration (progesterone and cetirizine). A third compound (RO-2) is likely to be similarly “overdosed” *in vitro*. And also, ascorbic acid gives positive signals only at clearly higher concentrations than usually found in human plasma. These considerations could be made more exactly on the basis of physiologically-based pharmacokinetic models and considerations of free drug concentrations. However, the principle is exemplified here, and if e.g., four of the false-positives would be negative at clinically-realistic concentrations, the test specificity would be 76%.

*TeraTox* offers additional benefits that are not yet available in any existing models. First, *TeraTox* is more sensitive than either the mEST or the *DevTox*^*qp*^ assay, especially when we consider maximum plasma concentrations (C_max_) from human data whenever possible or model species otherwise (Suppl. Tab. S2, S3). Detecting human-specific teratogens is critical for drug discovery and development, as illustrated by phthalimide-based series of molecules, which includes thalidomide (Smith and Mitchell 2018; Belair et al. 2020; Donovan et al. 2018; Matyskiela et al. 2018). Thalidomide was correctly identified as positive by *TeraTox*. Second, *TeraTox* reveals concentration-response teratogenicity and cytotoxicity relationship. This can be integrated with pharmacokinetic and exposure data to better estimate teratogenic risk in the clinic. Third, *TeraTox* generates quantitative gene expression data. Here we used this data to reveal germ-layer factors, to build a predictive model, and to identify pathways and gene-sets that are regulated by teratogens differently than by non-teratogens. Gene expression data can be also used to explore mechanisms of action and to prioritize drug candidates for preclinical development.

What sets *TeraTox* apart from other models is that it is less of a phenotypic black box but rather an interpretable and explainable model that provides mechanistic insights into gene, pathway, and germ-layer modulations. *TeraTox* informs predictions not only based on statistical data patterns but builds upon biological mechanisms and thus may reflect disturbed functionalities, similar to those leading to teratogenicity *in vivo*. These features put *TeraTox* conceptually in a group of other assays that use phenotypic changes or disturbed functionalities as readouts (Dreser et al. 2020; Hoelting et al. 2016; Meisig et al. 2020; Pallocca et al. 2016). The model consolidates our previous intention to ‘focus on germ layers’ and corroborates recent work exploring gastruloid models that profiles morphological changes of germ-layers for teratogenicity prediction (Jaklin et al. 2020; Moris et al. 2020).

Explainability analysis shed light on how the *TeraTox* model works and its limitations. Most importantly, we could distinguish cytotoxicity from teratogenicity. We explored machine-learning model variants for both teratogenicity and cytotoxicity predictions and made the observation that the best models depend on the target variable. Whereas germ-layer factors and random forest performed best for teratogenicity prediction, the combination of all pathway reporter genes and regularized linear regression with elastic nets showed the best prediction for cytotoxicity (Suppl. Fig. S3). There are two likely reasons. First, the molecular phenotyping platform contains well curated genes that reflect cytotoxicity and cell death, which were highlighted in a previous drug screening study using iPS-derived cardiomyocytes (Drawnel et al. 2017). Therefore, we anticipate that these genes are used by linear regression to predict cytotoxicity. Second, teratogenicity is notably complex. It can be induced in many different ways, with different perturbations leading to different down-stream changes that are collectively known as teratogenicity. Therefore, a change in the total output of the germ-layer regulatory network is probably a more robust readout of teratogenicity than individual genes. Random forest, which is an ensemble learning method, is better at detecting such heterogeneous signals than linear regression.

Further studies are warranted to explore several parallel paths for further optimization of the *TeraTox* assay. These can be divided into four categories: (1) paths leading to better characterization of EB differentiation, (2) paths leading to testing of larger chemical spaces beyond pharmaceuticals, (3) paths leading to better predictive and explanatory algorithms, and (4) paths leading to better biological models of human embryo development. These lines of research could broaden the applicability domain and increase the robustness of the *TeraTox* assay. To better characterize EBs, multi-modal characterizations of the EBs using bulk and single-cell omics, morphological profiling, and time-series experiments could be used. Extension of the assay duration to more than 7 days or using other differentiation protocols may further improve *TeraTox’* capacity to model mesoderm and endoderm development.

There are several options to further improve the predictivity and the explainability of the *TeraTox* model. To better distinguish between non-teratogens and teratogens, we may try to test the compounds with the *TeraTox* assay at lower concentrations (especially for non-teratogens), where the lowest concentration should be predicted to have a teratogenicity score equal to or close to zero. In this context, it could be feasible to apply the exposure-based validation approach described by Daston et al. (2014), based on minimal and maximal concentration-dependent effects of teratogenicity. Another option could be to include the *TeraTox* assay in a test battery to preselect those compounds that show high cytotoxic interference with weak teratogenicity. Multi-model data could be used to identify further relevant features beyond germ-layer genes and factors. As more data are collected, we may also optimize the prediction algorithm, for instance using the nearest-neighbor prediction or other variants. Finally, the *TeraTox* assay may benefit from a better modeling of human embryo development. We may use alternative morphology-based assays of gastruloids to complement the *TeraTox* readout (Baillie-Benson et al. 2020; Moris et al. 2020). Alternatively, sophisticated micro-physiological systems may better mimic the maternal-placenta-embryo axis and hence, may recapitulate true embryo exposure levels and give insights into active drug metabolism although drugs do not need to cross the placental barrier to cause fetal harm (Blundell et al. 2016; Blundell et al. 2018; Boos et al. 2021). In the future, they may replace the 3D embryoid bodies in *TeraTox*. In the current throughput, though, such systems will probably be more powerful as a secondary assay to spot check a few compounds of particular interest. For this purpose, a continuous integration and modeling of data of human embryogenesis, for instance from omics, imaging, and perturbation studies, is required to guide further optimization of the *TeraTox* assay (Canzler et al. 2020; Mantziou et al. 2021; Yan et al. 2013).

## 5. Conclusion

We demonstrate that the *TeraTox* is a novel predictive human *in vitro* assay for pharmaceutical teratogenicity prediction that addresses several limitations of other alternative assays regarding sensitivity, species-specificity, and explainability. We believe that its adoption in drug discovery empowers preclinical teratogenicity assessment. Further optimization of the *TeraTox* assay and its routine use in drug-screening processes will lead us towards better preclinical assessment of teratogenicity. Thus, we solicit the community for helping us with further refining and validating *TeraTox* in drug discovery and other contexts.

## 6. Acknowledgement

We thank Kevin Michaelsen, Claudia Bossen, and Jean-Christophe Hoflack for their generous support, as well as colleagues of the Bioinformatics and Exploratory Data Analysis (BEDA) team for their input and discussions.

## 7. Data availability

Sequencing files and gene counts are available at the NCBI GEO database (https://www.ncbi.nlm.nih.gov/geo/query/acc.cgi?acc=GSE183534). Differential gene expression data of commercially available compounds as well as the composition of germ-layer factors are available on the Zenodo platform (https://zenodo.org/record/6143691). All data are shared with the Creative Commons CC BY 4.0 license.

## References

Adams SS, Bough RG, Cliffe EE, Lessel B, Mills RFN. 1969. Absorption, distribution and toxicity of ibuprofen. Toxicology and Applied Pharmacology. 15(2):310–330.

Adler S, Pellizzer C, Hareng L, Hartung T, Bremer S. 2008. First steps in establishing a developmental toxicity test method based on human embryonic stem cells. Toxicol In Vitro. 22(1):200–211.

Augustyniak J, Bertero A, Coccini T, Baderna D, Buzanska L, Caloni F. 2019. Organoids are promising tools for species-specific in vitro toxicological studies. J Appl Toxicol.

Badillo S, Banfai B, Birzele F, Davydov II, Hutchinson L, Kam-Thong T, Siebourg-Polster J, Steiert B, Zhang JD. 2020. An introduction to machine learning. Clinical Pharmacology & Therapeutics. 107(4):871–885.

Baillie-Benson P, Moris N, Martinez Arias A. 2020. Pluripotent stem cell models of early mammalian development. Current Opinion in Cell Biology. 66:89–96.

Barredo Arrieta A, Díaz-Rodríguez N, Del Ser J, Bennetot A, Tabik S, Barbado A, Garcia S, Gil-Lopez S, Molina D, Benjamins R et al. 2020. Explainable artificial intelligence (xai): Concepts, taxonomies, opportunities and challenges toward responsible ai. Information Fusion. 58:82–115.

Barrow P. 2016. Revision of the ich guideline on detection of toxicity to reproduction for medicinal products: Swot analysis. Reproductive Toxicology. 64.

Beck F, Erler T, Russell A, James R. 1995. Expression of cdx-2 in the mouse embryo and placenta: Possible role in patterning of the extra-embryonic membranes. Dev Dyn. 204.

Belair DG, Lu G, Waller LE, Gustin JA, Collins ND, Kolaja KL. 2020. Thalidomide inhibits human ipsc mesendoderm differentiation by modulating crbn-dependent degradation of sall4. Sci Rep. 10(1):2864.

Blundell C, Tess ER, Schanzer AS, Coutifaris C, Su EJ, Parry S, Huh D. 2016. A microphysiological model of the human placental barrier. Lab Chip. 16(16):3065–3073.

Blundell C, Yi YS, Ma L, Tess ER, Farrell MJ, Georgescu A, Aleksunes LM, Huh D. 2018. Placental drug transport-on-a-chip: A microengineered in vitro model of transporter-mediated drug efflux in the human placental barrier. Adv Healthc Mater. 7(2).

Bock C, Kiskinis E, Verstappen G, Gu H, Boulting G, Smith ZD, Ziller M, Croft GF, Amoroso MW, Oakley DH et al. 2011. Reference maps of human es and ips cell variation enable high-throughput characterization of pluripotent cell lines. Cell. 144(3):439–452.

Boos JA, Misun PM, Brunoldi G, Furer LA, Aengenheister L, Modena M, Rousset N, Buerki-Thurnherr T, Hierlemann A. 2021. Microfluidic co-culture platform to recapitulate the maternal-placental-embryonic axis. Adv Biol (Weinh).e2100609.

Brannen KC, Chapin RE, Jacobs AC, Green ML. 2016. Alternative models of developmental and reproductive toxicity in pharmaceutical risk assessment and the 3rs. ILAR journal. 57(2):144–156.

Brannen KC, Charlap JH, Lewis EM. 2013. Zebrafish teratogenicity testing. In: Barrow PC, editor. Teratogenicity testing: Methods and protocols. Totowa, NJ: Humana Press.

Brooks M, Kristensen K, van Benthem K, Magnusson A, Berg CW, Nielsen A, Skaug H, Mächler M, Bolker B. 2017. Glmmtmb balances speed and flexibility among packages for zero-inflated generalized linear mixed modeling. R Journal. 9:378–400.

Burridge PW, Thompson S, Millrod MA, Weinberg S, Yuan X, Peters A, Mahairaki V, Koliatsos VE, Tung L, Zambidis ET. 2011. A universal system for highly efficient cardiac differentiation of human induced pluripotent stem cells that eliminates interline variability. PloS one. 6(4):e18293.

Canzler S, Schor J, Busch W, Schubert K, Rolle-Kampczyk UE, Seitz H, Kamp H, von Bergen M, Buesen R, Hackermüller J. 2020. Prospects and challenges of multi-omics data integration in toxicology. Archives of Toxicology.

Cassano A, Manganaro A, Martin T, Young D, Piclin N, Pintore M, Bigoni D, Benfenati E. 2010. Caesar models for developmental toxicity. Chem Cent J. 4 Suppl 1(Suppl 1):S4.

Chen JK, Taipale J, Cooper MK, Beachy PA. 2002. Inhibition of hedgehog signaling by direct binding of cyclopamine to smoothened. Genes & development. 16(21):2743–2748.

Cusack BJ, Parsons TE, Weinberg SM, Vieira AR, Szabo-Rogers HL. 2017. Growth factor signaling alters the morphology of the zebrafish ethmoid plate. Journal of Anatomy. 230(5):701–709.

Daniel S, Doron M, Fishman B, Koren G, Lunenfeld E, Levy A. 2019. The safety of amoxicillin and clavulanic acid use during the first trimester of pregnancy. British journal of clinical pharmacology. 85(12):2856–2863.

Dashe JS, Gilstrap LC, 3rd. 1997. Antibiotic use in pregnancy. Obstetrics and gynecology clinics of North America. 24(3):617–629.

Daston GP, Beyer BK, Carney EW, Chapin RE, Friedman JM, Piersma AH, Rogers JM, Scialli AR. 2014. Exposure-based validation list for developmental toxicity screening assays. Birth Defects Research Part B: Developmental and Reproductive Toxicology. 101(6):423–428.

Donovan KA, An J, Nowak RP, Yuan JC, Fink EC, Berry BC, Ebert BL, Fischer ES. 2018. Thalidomide promotes degradation of sall4, a transcription factor implicated in duane radial ray syndrome. Elife. 7.

Drawnel FM, Zhang JD, Küng E, Aoyama N, Benmansour F, Araujo Del Rosario A, Jensen Zoffmann S, Delobel F, Prummer M, Weibel F et al. 2017. Molecular phenotyping combines molecular information, biological relevance, and patient data to improve productivity of early drug discovery. Cell Chemical Biology. 24(5):624–634.e623.

Dreser N, Madjar K, Holzer AK, Kapitza M, Scholz C, Kranaster P, Gutbier S, Klima S, Kolb D, Dietz C et al. 2020. Development of a neural rosette formation assay (rofa) to identify neurodevelopmental toxicants and to characterize their transcriptome disturbances. Arch Toxicol. 94(1):151–171.

Etwel F, Djokanovic N, Moretti ME, Boskovic R, Martinovic J, Koren G. 2014. The fetal safety of cetirizine: An observational cohort study and meta-analysis. Journal of obstetrics and gynaecology : the journal of the Institute of Obstetrics and Gynaecology. 34(5):392–399.

Evans TJ. 2007. Chapter 14 - reproductive toxicity and endocrine disruption. In: Gupta RC, editor. Veterinary toxicology. Oxford: Academic Press. p. 206–244.

Genschow E, Spielmann H, Scholz G, Pohl I, Seiler A, Clemann N, Bremer S, Becker K. 2004. Validation of the embryonic stem cell test in the international ecvam validation study on three in vitro embryotoxicity tests. Alternatives to laboratory animals: ATLA. 32(3):209–244.

Genschow E, Scholz G, Brown N, Piersma A, Brady M, Clemann N, Huuskonen H, Paillard F, Bremer S, Becker K et al. 2000. Development of prediction models for three in vitro embryotoxicity tests in an ecvam validation study. In vitro & molecular toxicology. 13(1):51–66.

Hoelting L, Klima S, Karreman C, Grinberg M, Meisig J, Henry M, Rotshteyn T, Rahnenführer J, Blüthgen N, Sachinidis A et al. 2016. Stem cell-derived immature human dorsal root ganglia neurons to identify peripheral neurotoxicants. Stem Cells Transl Med. 5(4):476–487.

ICH. 2020. Ich harmonized guideline detection of reproductive and developmental toxicity for human pharmaceuticals s5(r3).

Jaklin M, Zhang JD, Barrow P, Ebeling M, Clemann N, Leist M, Kustermann S. 2020. Focus on germ-layer markers: A human stem cell-based model for in vitro teratogenicity testing. Reproductive Toxicology.

Kameoka S, Babiarz J, Kolaja K, Chiao E. 2013. A high-throughput screen for teratogens using human pluripotent stem cells. Toxicological Sciences. 137(1):76–90.

Krug AK, Kolde R, Gaspar JA, Rempel E, Balmer NV, Meganathan K, Vojnits K, Baquie M, Waldmann T, Ensenat-Waser R et al. 2013. Human embryonic stem cell-derived test systems for developmental neurotoxicity: A transcriptomics approach. Arch Toxicol. 87(1):123–143.

Lenz W, Knapp K. 1962. Foetal malformations due to thalidomide. Problems of birth defects. Springer. p. 200–206.

Lipinski RJ, Hutson PR, Hannam PW, Nydza RJ, Washington IM, Moore RW, Girdaukas GG, Peterson RE, Bushman W. 2008. Dose- and route-dependent teratogenicity, toxicity, and pharmacokinetic profiles of the hedgehog signaling antagonist cyclopamine in the mouse. Toxicol Sci. 104(1):189–197.

Ljosa V, Caie PD, ter Horst R, Sokolnicki KL, Jenkins EL, Daya S, Roberts ME, Jones TR, Singh S, Genovesio A et al. 2013. Comparison of methods for image-based profiling of cellular morphological responses to small-molecule treatment. Journal of Biomolecular Screening. 18(10):1321–1329.

Lüdecke D. 2018. Ggeffects: Tidy data frames of marginal effects from regression models. The Journal of Open Source Software. 3.

Mantziou V, Baillie-Benson P, Jaklin M, Kustermann S, Arias AM, Moris N. 2021. *in vitro* teratogenicity testing using a 3d, embryo-like gastruloid system. bioRxiv.2021.2003.2030.437698.

Marzo M, Kulkarni S, Manganaro A, Roncaglioni A, Wu S, Barton-Maclaren TS, Lester C, Benfenati E. 2016. Integrating in silico models to enhance predictivity for developmental toxicity. Toxicology. 370:127–137.

Matyskiela ME, Couto S, Zheng X, Lu G, Hui J, Stamp K, Drew C, Ren Y, Wang M, Carpenter A et al. 2018. Sall4 mediates teratogenicity as a thalidomide-dependent cereblon substrate. Nat Chem Biol. 14(10):981–987.

Meisig J, Dreser N, Kapitza M, Henry M, Rotshteyn T, Rahnenführer J, Hengstler JG, Sachinidis A, Waldmann T, Leist M et al. 2020. Kinetic modeling of stem cell transcriptome dynamics to identify regulatory modules of normal and disturbed neuroectodermal differentiation. Nucleic acids research. 48(22):12577–12592.

Moris N, Anlas K, van den Brink SC, Alemany A, Schröder J, Ghimire S, Balayo T, van Oudenaarden A, Martinez Arias A. 2020. An in vitro model of early anteroposterior organization during human development. Nature. 582(7812):410–415.

Muanda FT, Sheehy O, Bérard A. 2017. Use of antibiotics during pregnancy and the risk of major congenital malformations: A population based cohort study. British journal of clinical pharmacology. 83(11):2557–2571.

Pallocca G, Grinberg M, Henry M, Frickey T, Hengstler JG, Waldmann T, Sachinidis A, Rahnenführer J, Leist M. 2016. Identification of transcriptome signatures and biomarkers specific for potential developmental toxicants inhibiting human neural crest cell migration. Arch Toxicol. 90(1):159–180.

Palmer JA, Smith AM, Egnash LA, Colwell MR, Donley ELR, Kirchner FR, Burrier RE. 2017. A human induced pluripotent stem cell-based in vitro assay predicts developmental toxicity through a retinoic acid receptor-mediated pathway for a series of related retinoid analogues. Reprod Toxicol. 73:350–361.

Palmer JA, Smith AM, Egnash LA, Conard KR, West PR, Burrier RE, Donley EL, Kirchner FR. 2013. Establishment and assessment of a new human embryonic stem cell-based biomarker assay for developmental toxicity screening. Birth Defects Res B Dev Reprod Toxicol. 98(4):343–363.

Piersma AH. 1993. Whole embryo culture and toxicity testing. Toxicol In Vitro. 7(6):763–768.

Quintanilla RH, Jr., Asprer JST, Vaz C, Tanavde V, Lakshmipathy U. 2014. Cd44 is a negative cell surface marker for pluripotent stem cell identification during human fibroblast reprogramming. PloS one. 9(1):e85419.

Rempel E, Hoelting L, Waldmann T, Balmer NV, Schildknecht S, Grinberg M, Das Gaspar JA, Shinde V, Stober R, Marchan R et al. 2015. A transcriptome-based classifier to identify developmental toxicants by stem cell testing: Design, validation and optimization for histone deacetylase inhibitors. Arch Toxicol. 89(9):1599–1618.

Robinson MD, McCarthy DJ, Smyth GK. 2009. Edger: A bioconductor package for differential expression analysis of digital gene expression data. Bioinformatics. 26(1):139–140.

Rumbold A, Ota E, Nagata C, Shahrook S, Crowther CA. 2015. Vitamin c supplementation in pregnancy. Cochrane Database of Systematic Reviews. (9).

Sakata T, Chen JK. 2011. Chemical ‘jekyll and hyde’s: Small-molecule inhibitors of developmental signaling pathways. Chem Soc Rev. 40(8):4318–4331.

Scholz G, Genschow E, Pohl I, Bremer S, Paparella M, Raabe H, Southee J, Spielmann H. 1999a. Prevalidation of the embryonic stem cell test (est)-a new in vitro embryotoxicity test. Toxicol In Vitro. 13(4–5):675–681.

Scholz G, Pohl I, Genschow E, Klemm M, Spielmann H. 1999b. Embryotoxicity screening using embryonic stem cells in vitro: Correlation to in vivo teratogenicity. Cells, tissues, organs. 165(3–4):203–211.

Scutari M. 2010. Learning bayesian networks with the bnlearn r package. 2010. 35(3):22.

Seiler AE, Spielmann H. 2011. The validated embryonic stem cell test to predict embryotoxicity in vitro. Nat Protoc. 6(7):961–978.

Sergejew T. 2015. Evaluation of a human embryonic stem cell-based screening assay for the identification of teratogenic pharmaceuticals. THESIS FOR MASTER OF ADVANCED STUDIES IN TOXICOLOGY.

Shinde V, Hoelting L, Srinivasan SP, Meisig J, Meganathan K, Jagtap S, Grinberg M, Liebing J, Bluethgen N, Rahnenfuhrer J et al. 2017. Definition of transcriptome-based indices for quantitative characterization of chemically disturbed stem cell development: Introduction of the stop-toxukn and stop-toxukk tests. Arch Toxicol. 91(2):839–864.

Shinde V, Klima S, Sureshkumar PS, Meganathan K, Jagtap S, Rempel E, Rahnenfuhrer J, Hengstler JG, Waldmann T, Hescheler J et al. 2015. Human pluripotent stem cell based developmental toxicity assays for chemical safety screening and systems biology data generation. J Vis Exp. (100):e52333.

Shinde V, Perumal Srinivasan S, Henry M, Rotshteyn T, Hescheler J, Rahnenfuhrer J, Grinberg M, Meisig J, Bluthgen N, Waldmann T et al. 2016. Comparison of a teratogenic transcriptome-based predictive test based on human embryonic versus inducible pluripotent stem cells. Stem Cell Res Ther. 7(1):190.

Tsankov AM, Akopian V, Pop R, Chetty S, Gifford CA, Daheron L, Tsankova NM, Meissner A. 2015a. A qpcr scorecard quantifies the differentiation potential of human pluripotent stem cells. Nat Biotechnol. 33(11):1182–1192.

Tsankov AM, Gu H, Akopian V, Ziller MJ, Donaghey J, Amit I, Gnirke A, Meissner A. 2015b. Transcription factor binding dynamics during human es cell differentiation. Nature. 518(7539):344–349.

Vargesson N. 2015. Thalidomide-induced teratogenesis: History and mechanisms. Birth defects research Part C, Embryo today : reviews. 105(2):140–156.

Waldmann T, Grinberg M, König A, Rempel E, Schildknecht S, Henry M, Holzer A-K, Dreser N, Shinde V, Sachinidis A et al. 2016. Stem cell transcriptome responses and corresponding biomarkers that indicate the transition from adaptive responses to cytotoxicity. Chemical Research in Toxicology. 30(4):905–922.

Waldmann T, Rempel E, Balmer NV, Konig A, Kolde R, Gaspar JA, Henry M, Hescheler J, Sachinidis A, Rahnenfuhrer J et al. 2014. Design principles of concentration-dependent transcriptome deviations in drug-exposed differentiating stem cells. Chem Res Toxicol. 27(3):408–420.

Wang X, Moon J, Dodge ME, Pan X, Zhang L, Hanson JM, Tuladhar R, Ma Z, Shi H, Williams NS et al. 2013. The development of highly potent inhibitors for porcupine. Journal of medicinal chemistry. 56(6):2700–2704.

Whitlow S, Bürgin H, Clemann N. 2007. The embryonic stem cell test for the early selection of pharmaceutical compounds. ALTEX-Alternatives to animal experimentation. 24(1):3–7.

Worley KE, Rico-Varela J, Ho D, Wan LQ. 2018. Teratogen screening with human pluripotent stem cells. Integr Biol (Camb). 10(9):491–501.

Wu S, Fisher J, Naciff J, Laufersweiler M, Lester C, Daston G, Blackburn K. 2013. Framework for identifying chemicals with structural features associated with the potential to act as developmental or reproductive toxicants. Chem Res Toxicol. 26(12):1840–1861.

Yan L, Yang M, Guo H, Yang L, Wu J, Li R, Liu P, Lian Y, Zheng X, Yan J et al. 2013. Single-cell rna-seq profiling of human preimplantation embryos and embryonic stem cells. Nat Struct Mol Biol. 20(9):1131–1139.

Young DW, Bender A, Hoyt J, McWhinnie E, Chirn G-W, Tao CY, Tallarico JA, Labow M, Jenkins JL, Mitchison TJ et al. 2008. Integrating high-content screening and ligand-target prediction to identify mechanism of action. Nature Chemical Biology. 4(1):59–68.

Zhang JD, Hatje K, Sturm G, Broger C, Ebeling M, Burtin M, Terzi F, Pomposiello SI, Badi L. 2017. Detect tissue heterogeneity in gene expression data with bioqc. BMC Genomics. 18(1):277.

Zhang JD, Küng E, Boess F, Certa U, Ebeling M. 2015. Pathway reporter genes define molecular phenotypes of human cells. BMC Genomics. 16(1):342.

Zhang JD, Schindler T, Küng E, Ebeling M, Certa U. 2014. Highly sensitive amplicon-based transcript quantification by semiconductor sequencing. BMC Genomics. 15(1):565.

Zurlinden TJ, Saili KS, Rush N, Kothiya P, Judson RS, Houck KA, Hunter ES, Baker NC, Palmer JA, Thomas RS et al. 2020. Profiling the toxcast library with a pluripotent human (h9) stem cell line-based biomarker assay for developmental toxicity. Toxicological Sciences. 174(2):189–209.

