## Supplementary Data for "Optimization of the *TeraTox* assay for preclinical teratogenicity assessment"

### Supplementary information for Optimization of the TeraTox assay for preclinical teratogenicity assessment by Jaklin and Zhang et al.

#### Suppl. Materials and Methods

#### *Mouse Embryonic Stem Cell Test*

The test compounds used for the development of the *TeraTox* have also been tested in the mEST to compare the performance of both assays, with respect to investigate the strengths and limitations of the particular systems in terms of future strategic use of the assays. The protocol of the mEST was adapted from the original publication from Genschow *et al*., 2004 (Genschow et al. 2004) into an industry compliant format (Whitlow et al. 2007). We used the pluripotent mouse embryonic stem cell line ES-D3 (ATCC, CRL-1934) and the somatic mouse 3T3 fibroblast line (Balb/c 3T3 cell clone A31 from ATCC, CCL-163) and maintained them according to the ECVAM protocol. Most manual steps of the assay, such as cell seeding, dilution and addition of compounds, centrifugation and incubation of the EBs, are standardized and automated to gain reproducible data. The only non-automated assay procedures are cell maintenance and the manual count of beating cardiomyocytes.

The mEST assay is performed in two steps. First, the MTT cytotoxicity assay (3-(4,5-Dimethylthiazol-2-yl)-2,5-diphenyltetrazolium bromide) is conducted with both differentiated 3T3 fibroblasts and pluripotent D3 ESCs in monolayer cultures. Second, EBs derived from D3 ESCs are differentiated into cardiomyocytes over a total time course of 10 days, with compound treatment in six different concentrations on day 0, day 4 and day 7 in triplicates including 5-FU as positive reference. Test compounds and testing ranges with dilution ratios in 6 steps are shown in Suppl. Tab. S1. The concentration ranges were chosen based on cytotoxicity measurements in pilot experiments. Since the human and the murine test system differ in cellular origin, design and endpoints, the concentration ranges also differ between the assays. The endpoints measured are the concentration at which 50% inhibition of growth of 3T3 ($IC_{50}$ 3T3) and D3 cells ($IC_{50}$ D3) is achieved, and the concentration at which 50% inhibition of differentiation into cardiomyocytes ($ID_{50}$ D3) is achieved, compared to DMSO solvent controls, respectively (Suppl. Fig. S1a).

A modified discriminant function analysis was used to classify the test chemicals into two groups based on the calculated predictive score (PS) for a low potential of teratogenicity (negative, PS <0.6) and high potential of teratogenicity (positive, PS ≥0.6). This cut-off was implemented based on a previous internal validation, as a clear discrimination of possible teratogenic compounds is required in the drug development process. A possible prediction result is 'borderline' if calculated predictive scores are below the cut-off of 0.6 but above 0.5. Inconclusive results are also possible, for example, if solubility limits the concentration ranges tested to an extent that no $IC_{50}$ or $ID_{50}$ values can be reliably determined for one or more concentration-response curves (Suppl. Fig. S1b).

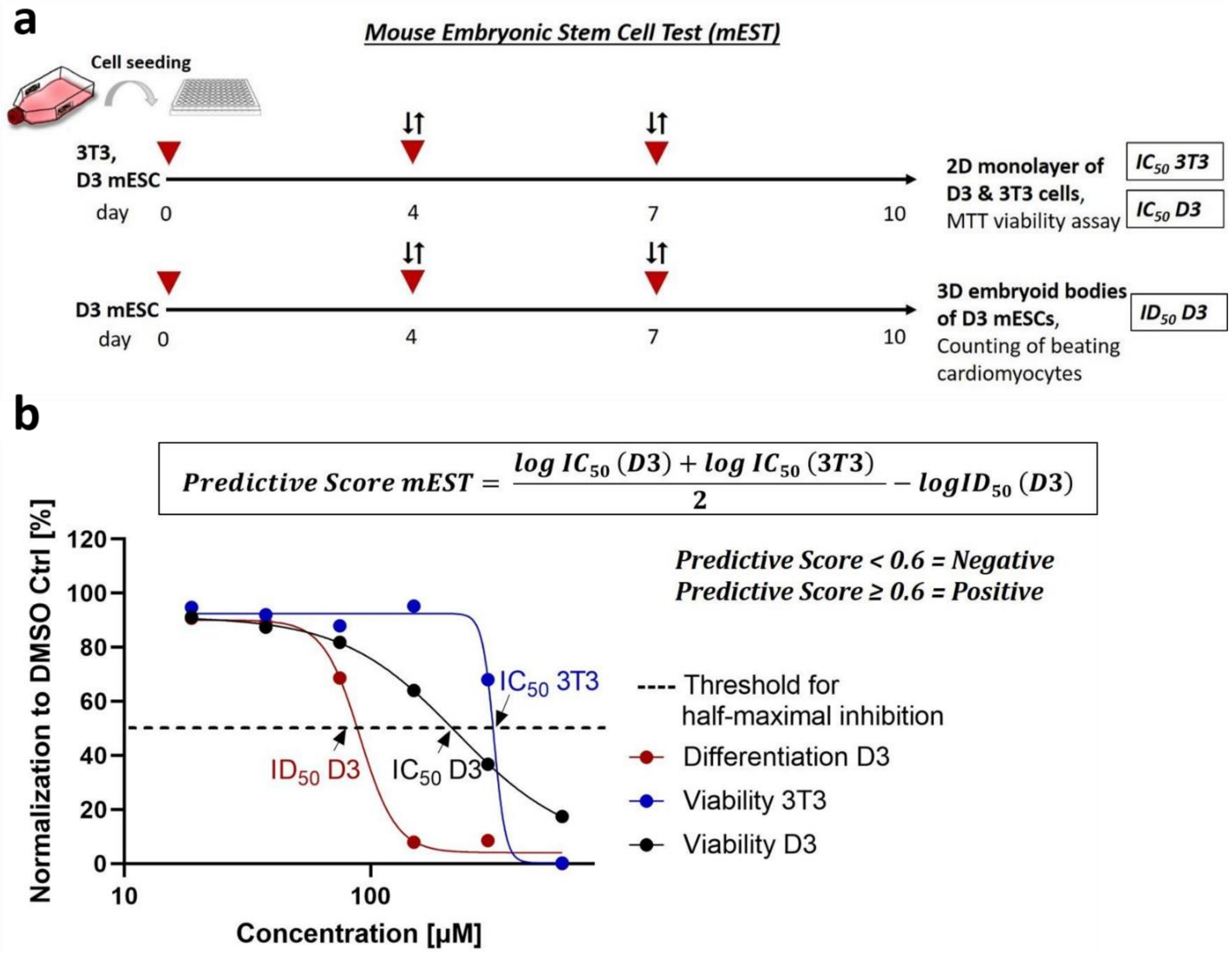


**Supplementary Figure *S*1: Mouse embryonic stem-cell test (mEST): assay workflow and prediction model.**

**(a)** Assay workflow. Mouse D3 and 3T3 cells are differentiated over a time course of 10 days to determine mEST endpoints ($IC_{50}$ and $ID_{50}$) for the calculation of the teratogenicity prediction score. Cells are treated with compounds in six concentrations at day 0, day 4, and day 7. The endpoints are cell viability (D3 and 3T3 cells) and counting of beating cardiomyocytes at day 10 (derived from D3 EBs).

**(b)** The prediction model of the mEST assay and normalized concentration-response curves. Values are normalized to solvent (DMSO) controls to determine the half-maximal concentration of viability for D3 ESC ($IC_{50}$ D3) and 3T3 fibroblasts ($IC_{50}$ 3T3) and half-maximal concentration of cardiomyocyte differentiation ($ID_{50}$ D3), respectively. A predictive score of <0.6 indicates non-teratogens whereas a predictive score ≥0.6 indicates teratogens.

**Suppl. Table S1: Reference compounds used in the mEST, with human teratogenicity classification and test concentration according to non-cytotoxic ranges (dilution ratios in brackets covering 6 concentrations).** Teratogenicity classification was based on FDA classification (Suppl. Tab. S2) or *in vivo* EFD data (indicated with asterisks*, Suppl. Tab. S3).

| ***Reference Compound*** | ***Teratogenicity Classification*** | ***Test Concentrations (mEST) [µM]*** |
|---|---|---|
| **Acitretin** | **Positive** | 0.02 – 100 (1:4) |
| **Amoxicillin** | Negative | 39 – 2500 (1:2) |
| **Artesunate** | **Positive** | 0.016 – 250 (1:6) |
| **Ascorbic Acid** | Negative | 0.043 – 2000 (1:6) |
| **Bosentan** | **Positive** | 7.8 – 500 (1:2) |
| **Busulfan** | **Positive** | 0.6 – 500 (1:3) |
| **Carbamazepine** | **Positive** | 11.7 – 750 (1:2) |
| **Cetirizine** | Negative | 11.7 – 750 (1:2) |
| **Cyclopamine** | **Positive** | 0.07 – 50 (1:3) |
| **Cyproheptadine** | Negative | 0.3 – 250 (1:3) |
| **Dabrafenib** | **Positive** | 0.14 – 100 (1:3) |
| **DAPT** | **Positive** | 0.12 – 500 (1:4) |
| **Dasatinib** | **Positive** | 0.3 – 20 (1:2) |
| **Dexamethasone** | **Positive** | 0.24 – 1000 (1:4) |
| **Dorsomorphin** | **Positive** | 0.01 – 50 (1:4) |
| **Doxycycline** | Negative | 2.0 – 1500 (1:3) |
| **5-Fluorouracil** | **Positive** | 0.03 – 20 (1:3) |
| **Hydroxyurea** | **Positive** | 7.8 – 500 (1:2) |
| **Ibuprofen** | Negative | 47 – 3000 (1:2) |
| **Isotretinoin** | **Positive** | 0.0001 – 250 (1:8) |
| **Imatinib** | **Positive** | 0.8 – 50 (1:2) |

| | | |
|---|---|---|
| **IWP-2** | **Positive** | 0.8 – 50 (1:2) |
| **Lazabemide** | Negative | 0.54 – 400 (1:3) |
| **Metformin** | Negative | 0.7 – 500 (1:3) |
| **Methotrexate** | **Positive** | 0.00006 – 1 (1:5) |
| **Misoprostol** | **Positive** | 1.6 – 100 (1:2) |
| **Penicillin G** | Negative | 31 – 2000 (1:2) |
| **Progesterone** | Negative | 0.7 – 500 (1:3) |
| **Retinoic Acid** | **Positive** | 0.0002 – 350 (1:11) |
| **RO-1*** | **Positive** | 4.7 – 300 (1:2) |
| **RO-2*** | Negative | 0.3 – 250 (1:3) |
| **RO-3*** | **Positive** | 7.8 – 500 (1:2) |
| **RO-4*** | Negative | 7.8 – 500 (1:2) |
| **RO-5*** | Negative | 0.07 – 50 (1:3) |
| **RO-6*** | Negative | 0.00025 – 250 (1:10) |
| **RO-7*** | Negative | 0.6 – 400 (1:2) |
| **RO-8*** | **Positive** | 1.9– 125 (1:2) |
| **RO-9*** | **Positive** | 0.8 – 50 (1:2) |
| **RO-10*** | **Positive** | 0.07 – 50 (1:3) |
| **RO-11*** | Negative | 2.3 – 150 (1:2) |
| **RO-12*** | Negative | 3.9 – 250 (1:2) |
| **SB431542** | **Positive** | 0.14 –100 (1:3) |
| **(±) Thalidomide** | **Positive** | 31.25 – 2000 (1:2) |
| **Valproic Acid** | **Positive** | 47 –3000 (1:2) |
| **Warfarin** | **Positive** | 39– 2500 (1:2) |

### *Factor analysis*

Factor analysis, sometimes called exploratory factor analysis to differentiate it from confirmatory factor analysis, is a statistical method to discover latent (unobserved) variables that account for the correlations observed between features. Useful for both dimension reduction and feature engineering, factor analysis has been particularly powerful in building predictive machine-learning models in biology using highly correlated features such as cell morphology in the context of high-content screening (Fabrigar and Wegener 2012; Hochreiter et al. 2006; Ljosa et al. 2013; Young et al. 2008).

With respect to gene expression, factor analysis reduces the data dimension from genes to factors, each of which is usually associated with multiple genes. Genes in each factor show correlated gene expression profiles across samples (Fig. 2a, b). These factors, therefore, can be thought of as being a representation of all biological processes influencing gene expression, for instance epigenetic profiles, transcription factor activities, microRNA abundances, etc. Despite the fact that most of these variables are not directly observable, latent factor analysis offers a possibility to infer their total contribution to detected variation in gene expression profiles.

Conceptually, factor analysis is familiar with other correlation-based methods, for instance Relevance Networks (BUTTE and KOHANE) and Weighted Correlation Network Analysis (WGCNA) (Langfelder and Horvath 2008). We preferred factor analysis to alternative methods because factor analysis does not make any additional assumptions than the common, minimum ones underlying correlation analyzes (homogeneity, completeness, etc.), whereas other methods do so, for instance the scale-free network structure assumed by WGCNA, whereas this assumption is often challenged (Broido and Clauset 2019; Khanin and Wit 2006). On the other hand, we have many more samples than the number of factors. Factor analysis is feasible with the maximum-likelihood method. We therefore decided to use factor analysis following the principle of Occam's Razor, *i.e.* prefer the simpler model unless more complex models are necessary.

#### *Teratogenicity Score*

A key challenge for building a predictive model of teratogenicity is that the potential of a compound inducing teratogenicity varies by its concentration. A concentration-response relationship can be assumed, namely a treatment with a higher concentration is more likely to induce teratogenicity than that with a low concentration. However, the concrete functional form between the potential and the concentration is not known. This motivated us to define the Teratogenicity Score as the '0-1 cosine bounded similarity' between differential gene expression profiles induced by any given concentration and the profiles induced by the maximum non-cytotoxic concentration.

Two important technical details require clarification. First is the range of the teratogenicity score. Mathematically, cosine similarity ranges between -1 and 1; we bounded it to 0-1 by setting negative similarities as zero, which did not change the performance of the models (data not shown) but helped with human understanding. The teratogenicity score can be interpreted as an estimate of the probability of inducing teratogenicity, which would be a real number between 0 and 1, though the real probability is unknown to us because we are working with an *in vitro* system only, and the probability estimated in our system may differ significantly from that *in vivo*.

The second technical detail is the selection of regression models. Given the truncated domain where the teratogenicity score is defined, we tried both simple linear regression and generalized linear models with beta regression. However, beta regression was computationally intensive and much slower, and its use led to similar results as simple linear regression for predicting teratogenicity scores. Therefore, we used simple linear regression throughout the study except in the last part of model explainability, because only one model is required there and the boundary consideration is important for simulation studies.

#### *Irregular concentration-response curves: causes and remedies*

Several reasons may cause irregular concentration-response curves deviating from the well-known form of a Hill function, for instance in the case of misoprostol and 5-FU. One reason is the non-linear response of differential gene expression. The other is interference by cytotoxicity. Another reason is the intrinsic property of a machine learning model: the prediction of the unseen test data was made on training data, and for the purpose of not introducing unnecessary biases, we did not specify the model to predict monotonic outcomes.

Fortunately, there are remedies against all these ill behaviors if they appear. First, we can investigate how gene expression responds to increasing concentrations both algorithmically, for instance with linear regression models, and visually, for instance with volcano plots. Second, we report both cytotoxicity and predicted TS in a visual form as in Figure 4C and in the supplementary figure. Last, we used various techniques, among others simulation studies (Figure 5d), to learn how the model makes its prediction. The results suggest that the model uses biological-relevant information, *i.e.* germ layers, for its prediction.

Other possible reasons include impact by outliers in the data, and/or technical errors and variabilities, which we have tried our best to avoid and exclude with optimal experiment design, quality control, and automation of the assay.

**Supplementary Equations**

$$Recall\ (sensitivity)\ =\ \frac{TP}{TP+FN} \qquad \text{(Eq. 1)}$$

$$Specificity\ = \frac{TN}{TN+FP} \qquad \text{(Eq. 2)}$$

$$Precision\ =\ \frac{TP}{TP+FP} \qquad \text{(Eq. 3)}$$

$$Accuracy\ =\ \frac{TP+TN}{TP+FP+TN+FN} \qquad \text{(Eq. 4)}$$

$$F_1 = \frac{2}{\frac{1}{Recall}+\frac{1}{Precision}} \qquad \text{(Eq. 5)}$$

**Supplementary Data**

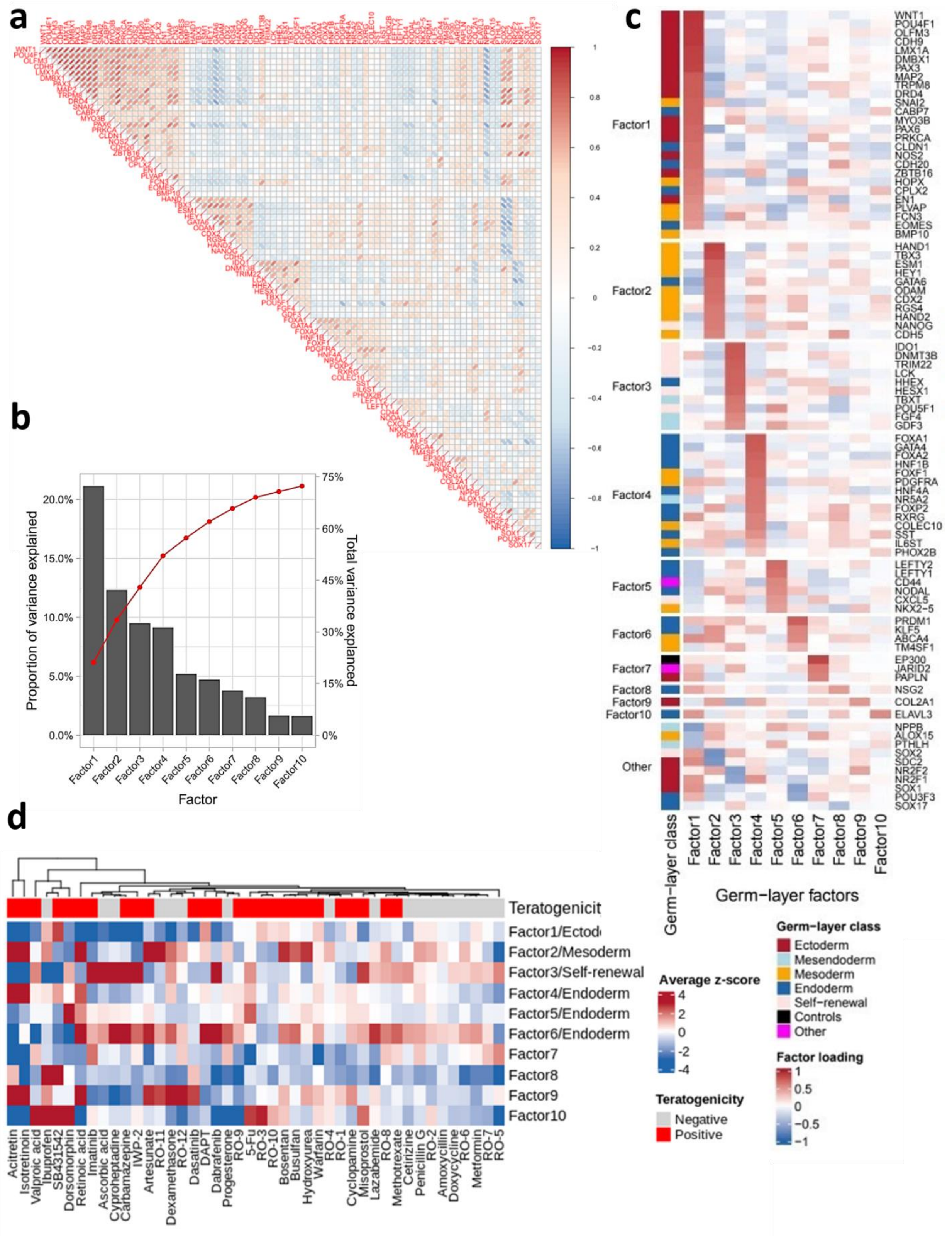

**Supplementary Figure S2: Factor analysis of germ-layer genes.**

**(a)** Pairwise Pearson correlation coefficients between germ-layer genes. Each row and each column indicate one gene. Red colors represent a strong correlation between expressions of genes whereas blue colors represent anti-correlation.

**(b)** Proportion of (co)variance explained by the first ten germ-layer factors (grey bars). Cumulatively, they explain more than 70% of total data variance (red line).

**(c)** Factor loading (like Figure 2b), with all gene symbols shown.

**(d)** Expression levels of germ-layer factors, represented as average z-scores of associated germ-layer genes, induced by compound treatments in the highest non-cytotoxic concentration (viability ≥ 80%). Blue colors indicate downregulation whereas red colors indicate upregulation of factors. The top side bar uses color to indicate compound classification: grey=non-teratogens, red=teratogens.

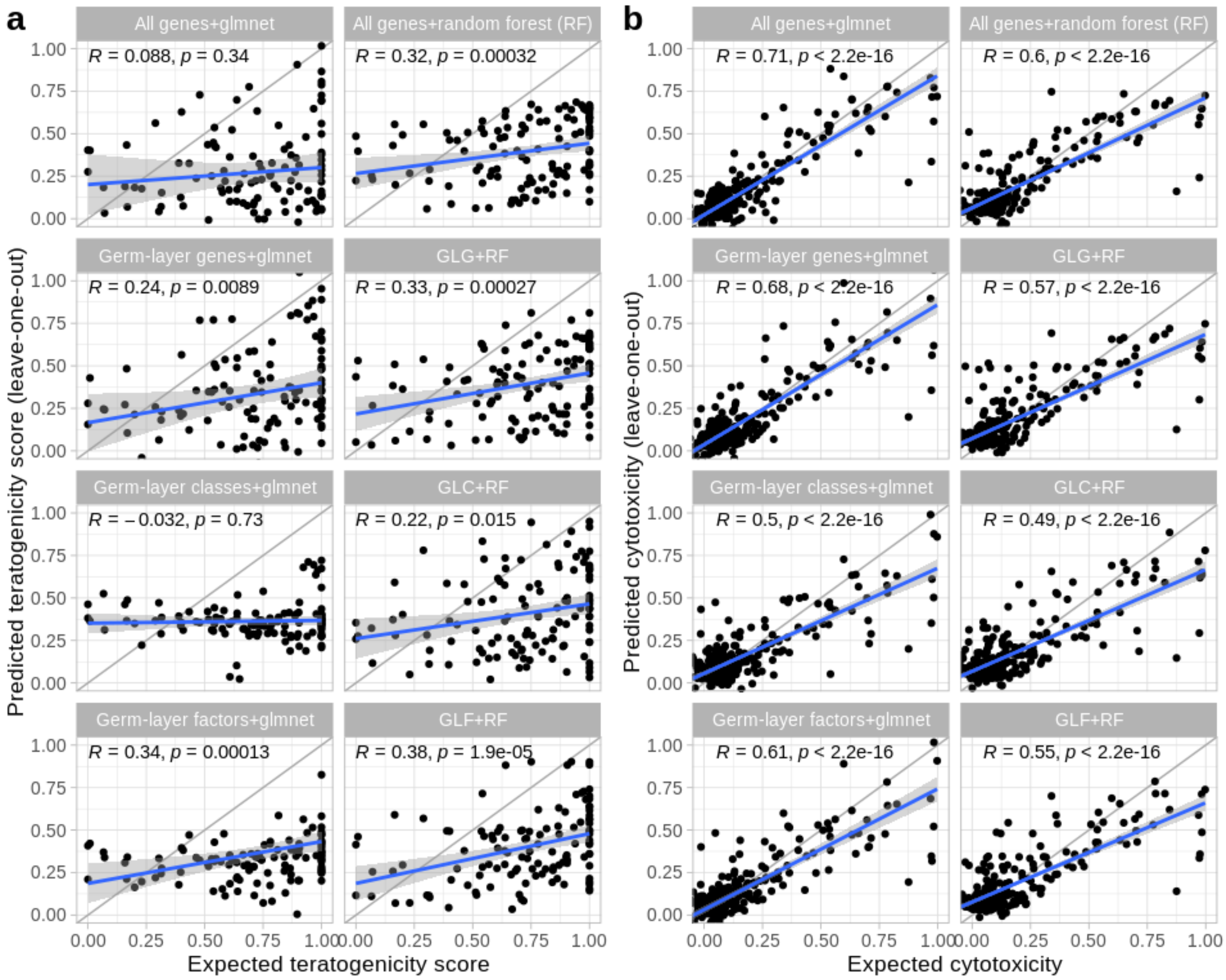


**Supplementary Figure S3: Leave-one-out prediction of teratogenicity scores and of cytotoxicity.**

**(a)** Prediction of teratogenicity scores using the leave-one-out scheme. For each combination of features and machine learning models, the teratogenicity score of each compound is predicted based on the scores of all other compounds. The predicted scores (y-axis) are compared with expected scores as defined in Fig. 3b. Each dot represents one concentration of one compound. The gray line indicates $y=x$. $R$ gives the Spearman correlation coefficient, and $p$ values are derived from the Spearman correlation test. The gray diagonal line represents $y=x$. The blue line indicates linear regression, with 95% confidence intervals in the gray area.

**(b)** Prediction of cytotoxicity using the leave-one-out scheme. All legends follow the definitions in Suppl. Fig. S3a.

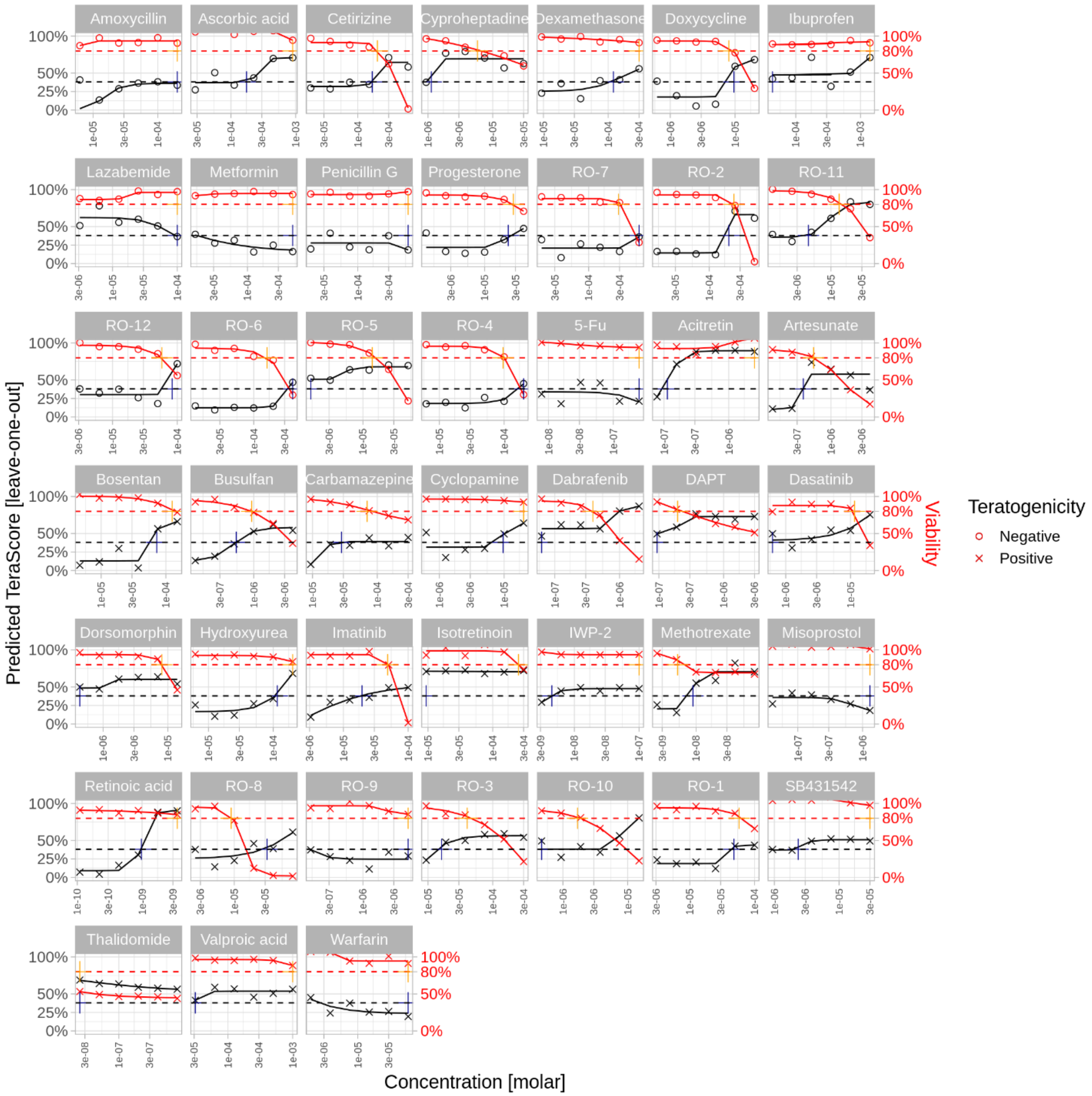


**Supplementary Figure S4: Classification of teratogenicity by Random Forest models.**

Concentration response curves of 45 reference compounds tested with the human *TeraTox* assay. Values for predicted teratogenicity were obtained by leave-one-out training/testing (median values of n=2), values for cytotoxicity were measured in the assay and normalized by DMSO (which is set as 100%). $NCC_{max}$ and $TC_{min}$ are plotted based on at least 80% viability threshold and the optimal teratogenicity score threshold of 0.38. Curve fit was performed with the four-parameter model offered by the *drc* package.

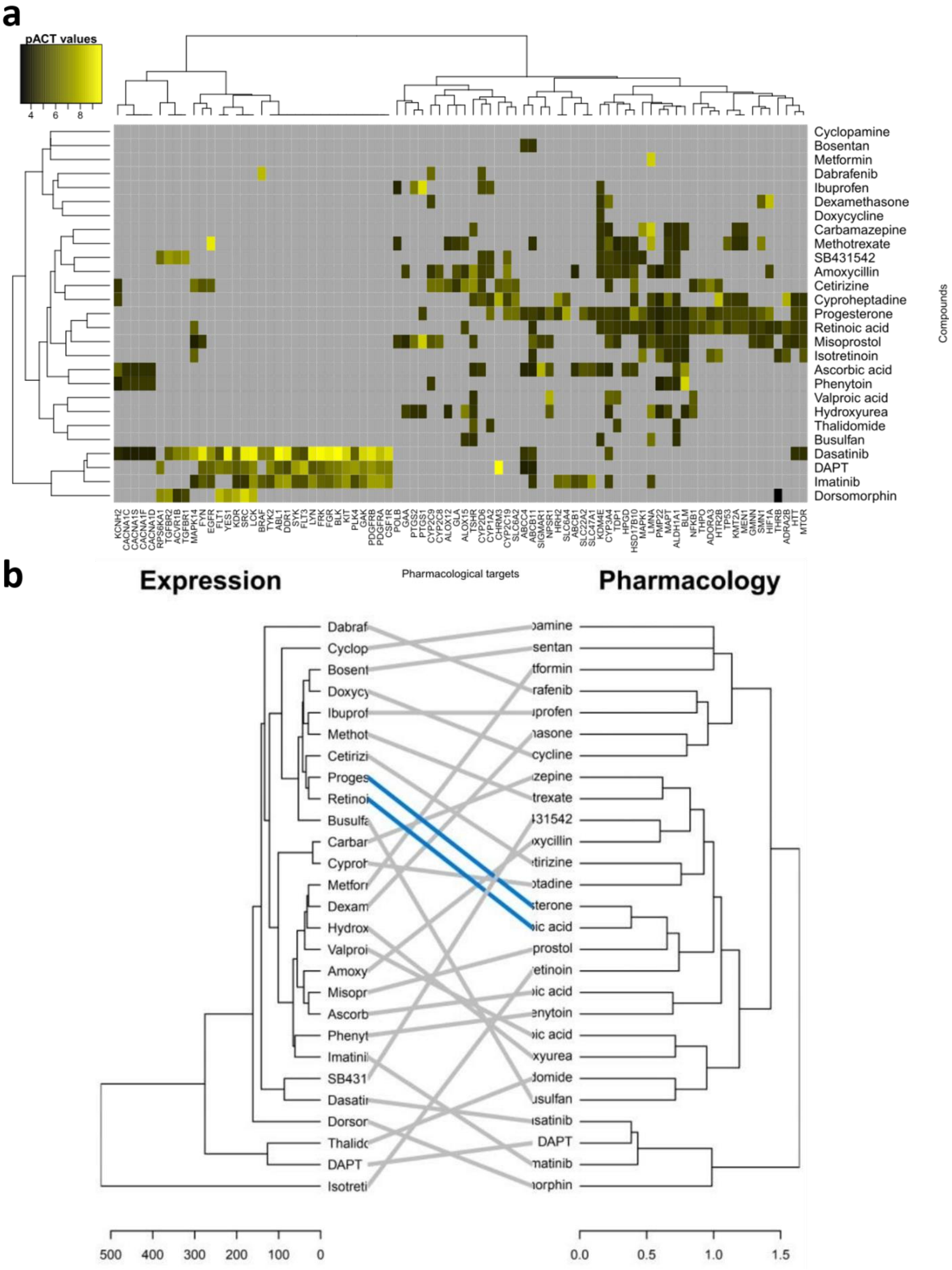
a
pACT values
4
6
8
Cyclopamine
Bosentan
Metformin
Dabrafenib
Ibuprofen
Dexamethasone
Doxycycline
Carbamazepine
Methotrexate
SB431542
Amoxycillin
Cetirizine
Cyproheptadine
Progesterone
Retinoic acid
Misoprostol
Isotretinoin
Ascorbic acid
Phenytoin
Valproic acid
Hydroxyurea
Thalidomide
Busulfan
Dasatinib
DAPT
Imatinib
Dorsomorphin
Compounds
KCNH2 CACNA1C CACNA1S CACNA1F CACNA1D RPS6KA1 TGFBR2 ACVR1B TGFBR1 MAPK14 FYN EGFR FLT1 YES1 KDR SRC LCK BRAF TYK2 ABL1 DDR1 SYK FLT3 LYN FRK FGR BLK KIT PLK4 GAK PDGFRB PDGFRA CSF1R POLB GAA PTGS2 PTGS1 CYP2C9 CYP2C8 ALOX12 GLA ALOX15 TSHR CYP2D6 CYP1A2 CHRM3 CYP2C19 SLC6A2 ABCC4 ABCB11 SIGMAR1 NPSR1 HRH2 SLC6A4 ABCB1 SLC22A2 SLC47A1 KDM4E CYP3A4 TDP1 HPGD HSD17B10 MAPK1 LMNA PMP22 MAPT ALDH1A1 BLM NFKB1 THPO ADORA3 HTR2B TP53 KMT2A MEN1 GMNN SMN1 HIF1A THRB ADRA2B HTT MTOR
Pharmacological targets
b
Expression
Pharmacology
500
400
300
200
100
0
0.0
0.5
1.0
1.5

**Supplementary Figure S5: Pharmacological profiles of the drugs.**

**(a)** Heatmap of target profiles of compounds. Only compounds with annotated targets in ChEMBL are shown. Each row represents one compound, and each column represents a human protein target. Colors indicate pACT values, which are absolute log10 transformed assay values (Ki, $IC_{50}$, *etc.*). Gray cells mean that data is not available. Compounds are clustered by binary distance and the Ward method.

**(b)** Two aligned dendrograms of differential gene expression and of pharmacology, linked by compounds. Left: dendrogram of differential gene expression profiles induced by compounds (average across concentration). Right: dendrogram of pharmacological profiles derived from Suppl. Fig. S5a. Lines connect the same compounds in two dendrograms. Gray lines: lack of correspondence. Blue lines: the same cluster is found in both dendrograms.

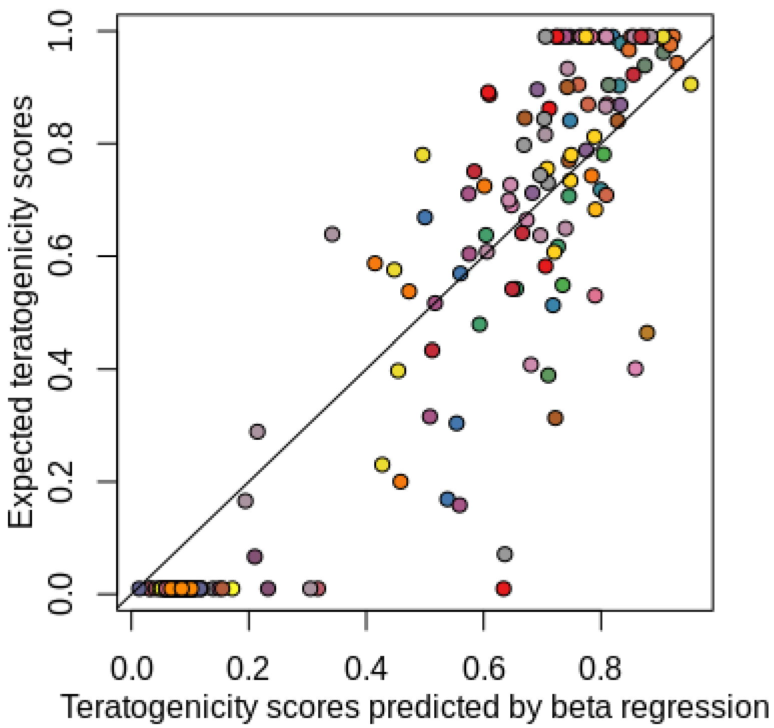


**Supplementary Figure S6: Performance of the generalized linear model with beta regression.**

Input to the regression model of ten germ-layer factors and significant interactions between them identified by a Bayesian network. The target variable is the teratogenicity score. We observe a good correlation between predicted (x-axis) and expected (y-axis) teratogenicity scores that are defined by cosine similarity. Each dot represents one concentration of one compound. Colors are used to represent different drugs. The diagonal line indicates *y*=*x*.

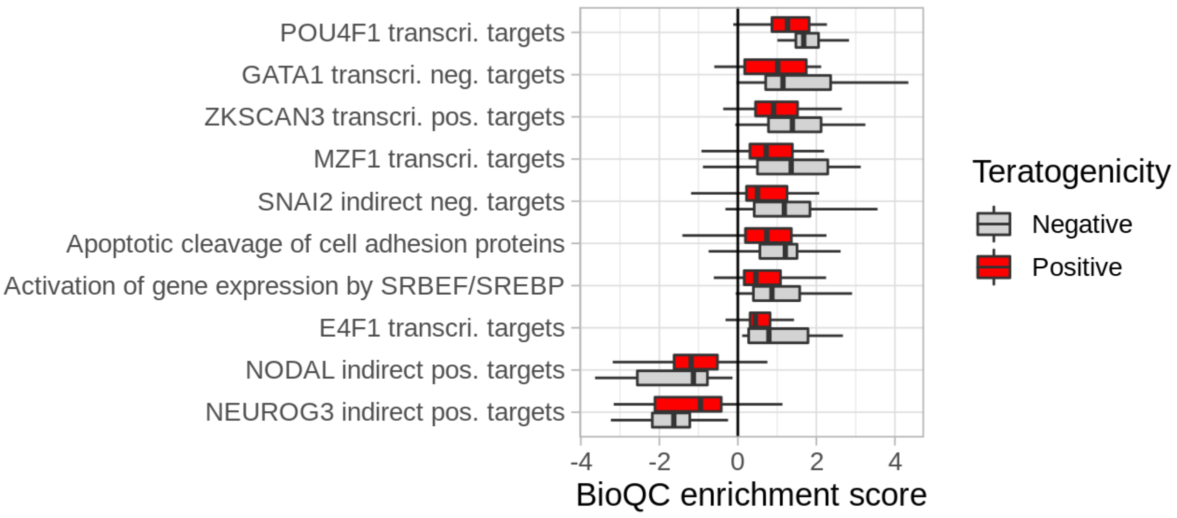


**Supplementary Figure S7: Different gene-set- and pathway-level regulation by teratogens and non-teratogens.**

We performed non-parametric Wilcoxon-Mann-Whitney tests on differential gene expression profiles of each tested compound using logFC as input, and derived a Q-score for each gene-set and each compound. The Q-score is absolute log10-transformed P-values of the two-sided Wilcoxon test. Then we applied a linear model to identify gene-sets with significantly different mean Q values between non-teratogens and teratogens. All gene-sets with P-value smaller than 0.10 are reported here in the boxplot. A negative Q-value indicates that the genes in the gene-set tend to be negatively regulated compared with DMSO, and a positive Q-value indicates that the genes tend to be positively regulated compared with DMSO. Abbreviations: transcri. = transcription; pos.= positive; neg.= negative.

**Supplementary Table S2: Human teratogenicity classification for commercial compounds** Classifications and human therapeutic plasma concentrations ($C_{max}$) were obtained from official data of the U.S. food and drug administration (FDA USPI) or from literature.

| ***Reference Compound*** | ***Teratogenicity Classification*** | ***Maximal therapeutic plasma concentration ($C_{max}$) [µM]*** | ***Reference*** |
|---|---|---|---|
| Acitretin | **Positive** | 2.4 | (FDA 2014; ICH 2020) |
| Amoxicillin | Negative | 13.8 | (Daniel et al. 2019; FDA 2019; Muanda et al. 2017) |
| Artesunate | **Positive** | 1.16 | (Amivas 2020; Morris et al. 2011) |
| Ascorbic Acid | Negative | 50 | (Martin et al. 1957; Mercer et al. 2010; Rumbold et al. 2015; Stephenson et al. 2013) |
| Bosentan | **Positive** | 1.1 | (Actelion Pharmaceuticals US 2008; ICH 2020; Li et al. 2018) |
| Busulfan | **Positive** | 0.52 | (GlaxoSmithKline 2003; ICH 2020) |
| Carbamazepine | **Positive** | 49 | (Corporation 2009; Güveli et al. 2017; Moreno et al. 2004; Tomson et al. 2019) |
| Cetirizine | Negative | 0.84 | (Etwel et al. 2014; ICH 2020; Pfizer 2002) |
| Cyclopamine | **Positive** | n/a | (Chen et al. 2002; Feutz and De Geyter 2019; Lipinski et al. 2008) |
| Cyproheptadine | Negative | 104.4 | (TevaPharmaceuticals 2014) |
| Dabrafenib | **Positive** | 2.8 | (GlaxoSmithKline 2014; ICH 2020) |

| | | | |
|---|---|---|---|
| DAPT | **Positive** | n/a | (Worley et al. 2018) |
| Dasatinib | **Positive** | 0.22 | (BMS 2010; ICH 2020; Rousselot et al. 2010) |
| Dexamethasone | **Positive** | 0.48 | (Merck & Co. 2019) |
| Dorsomorphin | **Positive** | n/a | (Sakata and Chen 2011) |
| Doxycycline | Negative | 13.6 | (FDA 2017; MaynePharma 2008; Muanda et al. 2017; Nahum et al. 2006; Newton et al. 2005) |
| 5-Fluorouracil | **Positive** | 222 | (Adler et al. 2008; ICH 2020) |
| Hydroxyurea | **Positive** | 684 | (Ballas et al. 2009; BMS 2016; Diav-Citrin et al. 1999; Gwer and Onyango 2018; ICH 2020; Thauvin-Robinet et al. 2001) |
| Ibuprofen | Negative | 286 | (Adams et al. 1969; Pfizer 2007) |
| Isotretinoin | **Positive** | 1 | (Dathe and Schaefer 2018; ICH 2020; Lammer et al. 1985; Nau 1993b; Piñeyro-Garza et al. 2015; Roche 2010) |
| Imatinib | **Positive** | 6.6 | (ICH 2020; Novartis 2012; von Mehren and Widmer 2011) |
| IWP-2 | **Positive** | n/a | (Kahn 2014; Qu et al. 2021; Warkus and Marikawa 2018) |
| Lazabemide | Negative | n/a | (Shepard and Lemire 2004) |

| | | | |
|---|---|---|---|
| Metformin | Negative | 9 | (BMS 2017; Montoya-Eguía et al. 2015) |
| Methotrexate | **Positive** | 4.7 | (Hyoun et al. 2012; ICH 2020; Kozma and Ramasethu 2011; Milunsky et al. 1968; Powell and Ekert 1971) |
| Misoprostol | **Positive** | n/a | (Kozma and Ramasethu 2011) |
| Penicillin G | Negative | 1150 | (Dashe and Gilstrap 1997; King Pharmaceuticals 2012; Muanda et al. 2017; Nahum et al. 2006; Wasz-Höckert et al. 1970) |
| Progesterone | Negative | 0.06 | (Ferring 2008; Paulson et al. 2014) |
| Retinoic Acid | **Positive** | 1.31 | (ICH 2020; Kam et al. 2012; Lammer et al. 1985; Nau 1993a; Shimozono et al. 2013) |
| SB431542 | **Positive** | n/a | (Belair et al. 2020) |
| (±) Thalidomide | **Positive** | 2.4 | (Belair et al. 2020; Celgene 2001; Donovan et al. 2018; ICH 2020; Matyskiela et al. 2018; Vargesson 2019) |
| Valproic Acid | **Positive** | 1423 | (Christensen et al. 2013; ICH 2020; Lloyd 2013; Tomson et al. 2019) |
| Warfarin | **Positive** | 6.8 | (FDA 1990; 2011; Walker et al. 2009) |

**Supplementary Table S3: *In vivo* EFD study data for developmental compounds.**

Developmental compounds provided by F. Hoffmann- La Roche (compound annotation was blinded due to confidential regulations). Studies for embryo-fetal development were either performed at Roche or contract research organizations. Compounds were generally classified as positive if effects were obtained in at least one species. Maximal protein plasma binding for the lowest observed adverse effect levels (LOAEL $C_{max}$) from the lowest dose where effects have been observed were averaged. Data were obtained from the Master Thesis of Thomas Sergejew 2015.

| ***Reference Compound*** | ***Teratogenicity Classification*** | ***in vivo data*** | | | |
|---|---|---|---|---|---|
| | | ***Rat*** | ***LOAEL $C_{max}$ [µM]*** | ***Rabbit*** | ***LOAEL $C_{max}$ [µM]*** |
| RO-1 | **Positive** | **Positive** | 18.5 | **Positive** | 37 |
| RO-2 | Negative | Negative | n/a | Negative | n/a |
| RO-3 | **Positive** | **Positive** | 91 | n/a | n/a |
| RO-4 | Negative | Negative | n/a | n/a | n/a |
| RO-5 | Negative | Negative | n/a | n/a | n/a |
| RO-6 | Negative | Negative | n/a | Negative | n/a |
| RO-7 | Negative | Negative | n/a | n/a | n/a |
| RO-8 | **Positive** | **Positive** | 54 | **Positive** | 32 |
| RO-9 | **Positive** | Negative | n/a | **Positive** | 0.94 |
| RO-10 | **Positive** | Negative | n/a | **Positive** | 41 |
| RO-11 | Negative | Negative | n/a | n/a | n/a |
| RO-12 | Negative | Negative | n/a | Negative | n/a |

**Supplementary Table S4: Germ-layer gene panel.**

Representative developmental markers (germ-layers) are classified into endoderm, ectoderm, mesoderm, mesendoderm, pluripotency (self-renewal) and other categories, as described by Tsankov *et al*. (Bock et al. 2011; Tsankov et al. 2015a; Tsankov et al. 2015b).

| Endoderm | Ectoderm | Mesoderm | Self-Renewal | Mesendoderm |
|---|---|---|---|---|
| CABP7 | WNT1 | SNAI2 | NANOG | T |
| CLDN1 | POU4F1 | HOPX | IDO1 | FGF4 |
| CDH20 | OLFM3 | PLVAP | DNMT3B | GDF3 |
| CPLX2 | CDH9 | FCN3 | TRIM22 | NR5A2 |
| EOMES | LMX1A | BMP10 | LCK | NPPB |
| GATA6 | DMBX1 | HAND1 | HESX1 | PTHLH |
| HHEX | PAX3 | TBX3 | POU5F1 | |
| FOXA1 | MAP2 | ESM1 | CXCL5 | |
| GATA4 | TRPM8 | HEY1 | SOX2 | |
| FOXA2 | DRD4 | ODAM | | |
| HNF1B | MYO3B | CDX2 | | |
| HNF4A | PAX6 | RGS4 | | |
| FOXP2 | PRKCA | HAND2 | | |
| RXRG | NOS2 | CDH5 | | |
| SST | ZBTB16 | FOXF1 | | |
| PHOX2B | EN1 | PDGFRA | | |
| LEFTY2 | PAPLN | COLEC10 | | |
| LEFTY1 | COL2A1 | IL6ST | | |
| NODAL | SDC2 | NKX2-5 | | |
| PRDM1 | NR2F2 | ABCA4 | | |
| KLF5 | NR2F1/NR2F2 | TM4SF1 | | |
| HMP19 | SOX1 | ALOX15 | | |
| ELAVL3 | | | | |
| POU3F3 | | | | |
| SOX17 | | | | |

**Supplementary Table S5: Teratogenicity Prediction of Reference Compounds**.

Classification of positive and negative reference compounds by the human *TeraTox* assay with associated maximum non-cytotoxic concentrations ($NCC_{max}$) and minimal teratogenic concentrations ($TC_{min}$) and its calculated predictive *TeraTox* score. Classification of the mouse EST with associated half-maximal concentrations for inhibition of growth for D3 mouse ESC ($IC_{50}D3$) and 3T3 fibroblasts ($IC_{50}3T3$) and half-maximal concentrations for inhibition of differentiation into beating cardiomyocytes for D3 mouse ESC ($ID_{50}D3$) and its calculated predictive score. TN= true negative, TP= true positive, FN= false negative, FP= false positive, values of several assay runs were averaged, n≥3, *highly cytotoxic

| ***Reference Compound*** | ***Teratogenicity classification*** | **Human *TeraTox* Assay** | | | | **Mouse embryonic stem cell test** | | | | |
|---|---|---|---|---|---|---|---|---|---|---|
| | | *$NCC_{max}$ [µM]* | *$TC_{min}$ [µM]* | *TeraTox score* | *Predicted Teratogenicity* | *$IC_{50}D3$ [µM]* | *$IC_{50}3T3$ [µM]* | *$ID_{50}D3$ [µM]* | *Predictive Score* | *Predicted Teratogenicity* |
| Amoxicillin | Negative | 200 | 200 | 0.00 | TN | 2500 | 2500 | 2467 | 0.01 | TN |
| Ascorbic acid | Negative | 900 | 174 | 0.71 | **FP** | 1104 | 2000 | 2000 | n/a | TN |
| Cetirizine | Negative | 201 | 167 | 0.08 | **FP** | 332 | 500 | 215 | 0.27 | TN |
| Cyproheptadine | Negative | 5.8 | 1.1 | 0.72 | **FP** | 12 | 45 | 0.7 | 1.55 | **FP** |
| Doxycycline | Negative | 8.0 | 9.6 | -0.08 | TN | 244 | 397 | 11.3 | 1.44 | **FP** |
| Ibuprofen | Negative | 1400 | 43.7 | 1.50 | **FP** | 2628 | 1380 | 1166 | 0.21 | TN |
| Lazabemide | Negative | 100 | 100 | 0.00 | TN | 59 | 235 | 106 | 0.05 | TN |
| Metformin | Negative | 500 | 500 | 0.00 | TN | 500 | 500 | 500 | n/a | TN |
| Penicillin G | Negative | 600 | 600 | 0.00 | TN | 2000 | 2000 | 2000 | n/a | TN |
| Progesterone | Negative | 28 | 23 | 0.08 | **FP** | 58 | 36 | 24 | 0.29 | TN |
| RO-7 | Negative | 289 | 600 | -0.32 | TN | 227 | 227 | 129 | 0.25 | TN |

| | | | | | | | | | | |
|---|---|---|---|---|---|---|---|---|---|---|
| RO-2 | Negative | 241 | 201 | 0.08 | **FP** | 47 | 68 | 90 | n/a | TN |
| RO-11 | Negative | 13 | 4.5 | 0.46 | **FP** | 32 | 13 | 5.3 | 0.59 | **BL** |
| RO-12 | Negative | 58 | 83 | -0.16 | TN | 92 | 137 | 37 | 0.48 | TN |
| RO-6 | Negative | 161 | 400 | -0.40 | TN | 386 | 515 | 438 | 0.01 | TN |
| RO-5 | Negative | 14 | 1.6 | 0.94 | **FP** | 9.1 | 5.8 | 4.1 | 0.25 | TN |
| RO-4 | Negative | 96 | 200 | -0.32 | TN | 39 | 170 | 2.5 | 1.51 | **FP** |
| 5-FU | **Positive** | 0.3 | 0.3 | 0.00 | **FN** | 0.5 | 2.0 | 0.3 | 0.48 | **FN** |
| Acitretin | **Positive** | 2.5 | 0.1 | 1.39 | TP | 0.2 | 129 | 0.0 | 2.64 | TP |
| Artesunate | **Positive** | 0.5 | 0.4 | 0.09 | TP | 3.2 | 6.5 | 1.3 | 0.53 | **FN** |
| Bosentan | **Positive** | 125 | 72 | 0.24 | TP | 19.3 | 70.6 | 22.1 | 0.22 | **FN** |
| Busulfan | **Positive** | 0.9 | 0.5 | 0.25 | TP | 19.3 | 70.6 | 22.1 | 0.22 | **FN** |
| Carbamazepine | **Positive** | 70 | 28 | 0.40 | TP | 372 | 393 | 207 | 0.26 | **FN** |
| Cyclopamine | **Positive** | 20 | 9.6 | 0.32 | TP | 22 | 76 | 5.9 | 0.84 | TP |
| Dabrafenib | **Positive** | 0.4 | 0.06 | 0.82 | TP | 23 | 23 | 20 | 0.07 | **FN** |
| DAPT | **Positive** | 0.2 | 0.09 | 0.35 | TP | 324 | 176 | 35 | 0.83 | TP |
| Dasatinib | **Positive** | 12 | 0.6 | 1.31 | TP | 4.3 | 0.6 | 3.7 | n/a | TP |
| Dexamethasone | **Positive** | 300 | 120 | 0.40 | TP | 87 | 300 | 55 | 0.46 | **FN** |
| Dorsomorphin | **Positive** | 8.1 | 0.4 | 1.31 | TP | 1.9 | 1.4 | 0.7 | 0.34 | **FN** |

| | | | | | | | | | | |
|---|---|---|---|---|---|---|---|---|---|---|
| Hydroxyurea | **Positive** | 200 | 116 | 0.24 | TP | 51 | 89 | 23 | 0.47 | **FN** |
| Imatinib | **Positive** | 48 | 19 | 0.40 | TP | 12 | 22 | 7 | 0.39 | **FN** |
| Isotretinoin | **Positive** | 250 | 9.4 | 1.42 | TP | 33 | 121 | 0.5 | 2.10 | TP |
| IWP-2 | **Positive** | 0.1 | 4.5E3 | 1.35 | TP | 15.5 | 55 | 1 | 1.47 | TP |
| Methotrexate | **Positive** | 5.2E3 | 9E3 | -0.23 | **FN** | 0.2 | 0.1 | 0.1 | 0.10 | **FN** |
| Misoprostol | **Positive** | 1.30 | 1.30 | 0.00 | **FN** | 23 | 13 | 40 | n/a | **FN** |
| Retinoic acid | **Positive** | 3.5E3 | 9.8E4 | 0.55 | TP | 0.004 | 77.9 | 0.014 | 1.60 | TP |
| RO-8 | **Positive** | 9.0 | 32 | -0.55 | **FN** | 110 | 104 | 85 | 0.10 | **FN** |
| RO-9 | **Positive** | 5.0 | 5.0 | 0.00 | **FN** | 40 | 21 | 3.7 | 0.89 | TP |
| RO-3 | **Positive** | 40 | 16 | 0.40 | TP | 77 | 147 | 18 | 0.77 | TP |
| RO-10 | **Positive** | 1.7 | 0.5 | 0.53 | TP | 5.1 | 4.2 | 1 | 0.67 | TP |
| RO1 | **Positive** | 58 | 48 | 0.08 | TP | 68 | 180 | 24 | 0.66 | TP |
| SB431542 | **Positive** | 30 | 2.3 | 1.11 | TP | 36 | 21 | 5.8 | 0.68 | TP |
| Thalidomide | **Positive** | 0.03 | 0.03 | n/a* | TP | 2000 | 2000 | 2000 | n/a | **FN** |
| Valproic acid | **Positive** | 1000 | 31 | 1.51 | TP | 1252 | 2859 | 441 | 0.63 | TP |
| Warfarin | **Positive** | 60 | 60 | 0.00 | **FN** | 1892 | 895 | 974 | 0.13 | **FN** |

**Supplementary Table S6: Teratogenicity Prediction of Reference Compounds used within TeraTox and DevTox assay by Stemina (ToxCast)**.

Classification of positive and negative reference compounds that have been measured in both assays, the human *TeraTox* assay in comparison to the DevTox$^{qp}$ assay by Stemina (ToxCast assessment). TN= true negative, TP= true positive, FN= false negative, FP= false positive.

| **Reference compound** | **Human *TeraTox* Assay** | **Stemina devTOX$^{qP}$** |
|---|---|---|
| All-trans Retinoic acid | TP | TP |
| Methotrexate | FN | TP |
| Thalidomide | TP | TP |
| 5-Fluorouracil | FN | TP |
| Carbamazepine | TP | TP |
| Busulfan | TP | TP |
| Dexamethasone | TP | TP |
| Hydroxyurea | TP | TP |
| Valproic Acid | TP | TP |
| Cyclopamine | TP | FN |
| Warfarin | FN | FN |
| Penicillin G | TN | TN |
| Bosentan | TP | FN |
| Ascorbic acid | FP | TN |
| Artesunate | TP | FN |
| Amoxicillin | TN | TN |
| Isotretinoin | TP | TP |
| Acitretin | TP | TP |

**Supplementary Table S7: Comparison of performance measures of the human TeraTox and DevTox$^{qp}$ assay by Stemina**.

Values were calculated based on a subset of 18 compounds based on the ToxCast compound set used within both assays, see Suppl. Tab. S6 (according to supplementary equations 1-5. TP= true positive, TN=true negative, FP=false positive, FN=false negative).

| ***Model*** | ***TP*** | ***TN*** | ***FP*** | ***FN*** | ***Accuracy*** | ***Balanced Accuracy*** | ***Precision*** | ***Recall*** | ***Specificity*** | ***$F_1$*** |
|---|---|---|---|---|---|---|---|---|---|---|
| ***DevTox$^{qp}$*** | 11 | 3 | 0 | 4 | 78% | 87% | 100% | 73% | 100% | 85% |
| ***TeraTox*** | 12 | 2 | 1 | 3 | 78% | 73% | 92% | 80% | 67% | 86% |

**Supplementary Table S8:** **QSAR/TeraTox/mEST comparisons, including 20 compounds**

| Reference Compound | Teratogenicity Classification | TeraTox Prediction | mEST Prediction | CAESAR Assessment | PG Assessment | CAESAR Reliability | PG Reliability |
|---|---|---|---|---|---|---|---|
| Acitretin | Positive | TP | TP | Toxicant | Developmental toxicant | GOOD reliability | EXPERIMENTAL value |
| Artesunate | Positive | TP | FN | Toxicant | NON-Toxicant | LOW reliability | LOW reliability |
| Bosentan | Positive | TP | FN | Toxicant | NON-Toxicant | LOW reliability | LOW reliability |
| Busulfan | Positive | TP | FN | Toxicant | NON-Toxicant | LOW reliability | LOW reliability |
| Carbamazepine | Positive | TP | FN | Toxicant | Reproductive and developmental toxicant | GOOD reliability | EXPERIMENTAL value |
| Cetirizine | Negative | FP | TN | Toxicant | Toxicant | MODERATE reliability | GOOD reliability |
| Cyclopamine | Positive | TP | TP | Toxicant | Developmental toxicant | MODERATE reliability | EXPERIMENTAL value |
| Dabrafenib | Positive | TP | FN | Toxicant | NON-Toxicant | LOW reliability | LOW reliability |
| DAPT | Positive | TP | TP | NON-Toxicant | NON-Toxicant | MODERATE reliability | LOW reliability |
| Dasatinib | Positive | TP | TP | Toxicant | NON-Toxicant | MODERATE reliability | LOW reliability |
| Dorsomorphin | Positive | TP | FN | Toxicant | NON-Toxicant | LOW reliability | LOW reliability |
| Doxycycline | Negative | TN | FP | Toxicant | Toxicant | GOOD reliability | GOOD reliability |
| Imatinib | Positive | TP | FN | Toxicant | NON-Toxicant | MODERATE reliability | LOW reliability |
| IWP-2 | Positive | TP | TP | NON-Toxicant | NON-Toxicant | LOW reliability | LOW reliability |
| Lazabemide | Negative | TN | TN | NON-Toxicant | NON-Toxicant | MODERATE reliability | LOW reliability |
| Metformin | Negative | TN | TN | NON-Toxicant | NON-Toxicant | LOW reliability | LOW reliability |
| Misoprostol | Positive | FN | FN | Toxicant | Reproductive and developmental toxicant | MODERATE reliability | EXPERIMENTAL value |
| Progesterone | Negative | FP | TN | Toxicant | Reproductive and developmental toxicant | GOOD reliability | EXPERIMENTAL value |
| SB431542 | Positive | TP | TP | Toxicant | NON-Toxicant | LOW reliability | LOW reliability |
| (±) Thalidomide | Positive | TP | FN | Toxicant | Developmental toxicant | GOOD reliability | EXPERIMENTAL value |

**Supplementary Table S9: Cost calculation of the human TeraTox and the mEST assay**

Values were calculated based on the consumables used for six compounds within both assays. Sequencing data not included (~50 USD/ sample). Values may vary in terms of different pricing conditions.

| **Cost factor** | **mEST** | **TeraTox** |
|---|---|---|
| Price per assay (based on 6 compounds) | 1,200 USD | 1,300 USD |
| Price per compound | 200 USD | 217 USD |
| Cell maintenance cost per week | 290 USD | 230 USD |
| Lab time effort | 20h | 12h |
| Assay duration | 10 days | 7 days (without sequencing) |
| max. throughput per assay | 6 compounds | more than 12 compounds |

### Supplementary References

Actelion Pharmaceuticals US I. 2008. Tracleer® [bosentan] https://wwwaccessdatafdagov/drugsatfda_docs/label/2009/021290s015lblpdf.

Adams SS, Bough RG, Cliffe EE, Lessel B, Mills RFN. 1969. Absorption, distribution and toxicity of ibuprofen. Toxicology and Applied Pharmacology. 15(2):310-330.

Adler S, Pellizzer C, Hareng L, Hartung T, Bremer S. 2008. First steps in establishing a developmental toxicity test method based on human embryonic stem cells. Toxicol In Vitro. 22(1):200-211.

Amivas. 2020. Artesunate for injection, for intravenous use. https://wwwaccessdatafdagov/drugsatfda_docs/label/2020/213036s000lblpdf.

Ballas SK, McCarthy WF, Guo N, DeCastro L, Bellevue R, Barton BA, Waclawiw MA. 2009. Exposure to hydroxyurea and pregnancy outcomes in patients with sickle cell anemia. Journal of the National Medical Association. 101(10):1046-1051.

Belair DG, Lu G, Waller LE, Gustin JA, Collins ND, Kolaja KL. 2020. Thalidomide inhibits human ipsc mesendoderm differentiation by modulating crbn-dependent degradation of sall4. Sci Rep. 10(1):2864.

BMS. 2010. Sprycel™ (dasatinib). https://wwwaccessdatafdagov/drugsatfda_docs/label/2010/021986s7s8lblpdf.

BMS. 2016. Hydrea (hydroxyurea) highlights of prescribing information https://wwwaccessdatafdagov/drugsatfda_docs/label/2016/016295Orig1s047,s048Lblpdf.

BMS. 2017. Glucophage® (metformin hydrochloride) tablets. https://wwwaccessdatafdagov/drugsatfda_docs/label/2017/020357s037s039,021202s021s023lblpdf.

Bock C, Kiskinis E, Verstappen G, Gu H, Boulting G, Smith ZD, Ziller M, Croft GF, Amoroso MW, Oakley DH et al. 2011. Reference maps of human es and ips cell variation enable high-throughput characterization of pluripotent cell lines. Cell. 144(3):439-452.

Broido AD, Clauset A. 2019. Scale-free networks are rare. Nature Communications. 10(1):1017.

BUTTE AJ, KOHANE IS. Mutual information relevance networks: Functional genomic clustering using pairwise entropy measurements. Biocomputing 2000. p. 418-429.

Celgene. 2001. Thalomid® capsules (thalidomide). https://wwwaccessdatafdagov/drugsatfda_docs/label/2001/20785s12s14lblpdf.

Chen JK, Taipale J, Cooper MK, Beachy PA. 2002. Inhibition of hedgehog signaling by direct binding of cyclopamine to smoothened. Genes & development. 16(21):2743-2748.

Christensen J, Grønborg TK, Sørensen MJ, Schendel D, Parner ET, Pedersen LH, Vestergaard M. 2013. Prenatal valproate exposure and risk of autism spectrum disorders and childhood autism. JAMA. 309(16):1696-1703.

Corporation NP. 2009. Tegretol® carbamazepine usp https://wwwaccessdatafdagov/drugsatfda_docs/label/2009/016608s101,018281s048lblpdf.

Daniel S, Doron M, Fishman B, Koren G, Lunenfeld E, Levy A. 2019. The safety of amoxicillin and clavulanic acid use during the first trimester of pregnancy. British journal of clinical pharmacology. 85(12):2856-2863.

Dashe JS, Gilstrap LC, 3rd. 1997. Antibiotic use in pregnancy. Obstetrics and gynecology clinics of North America. 24(3):617-629.

Dathe K, Schaefer C. 2018. Drug safety in pregnancy: The german embryotox institute. Eur J Clin Pharmacol. 74(2):171-179.

Diav-Citrin O, Hunnisett L, Sher GD, Koren G. 1999. Hydroxyurea use during pregnancy: A case report in sickle cell disease and review of the literature. American journal of hematology. 60(2):148-150.

Donovan KA, An J, Nowak RP, Yuan JC, Fink EC, Berry BC, Ebert BL, Fischer ES. 2018. Thalidomide promotes degradation of sall4, a transcription factor implicated in duane radial ray syndrome. Elife. 7.

Etwel F, Djokanovic N, Moretti ME, Boskovic R, Martinovic J, Koren G. 2014. The fetal safety of cetirizine: An observational cohort study and meta-analysis. Journal of obstetrics and gynaecology: the journal of the Institute of Obstetrics and Gynaecology. 34(5):392-399.

Fabrigar LR, Wegener DT. 2012. Exploratory factor analysis. Oxford: Oxford University Press.

FDA. 1990. Warfarin. https://wwwpharmapendiumcom/browse/fda/Warfarin%20Sodium/b223ffd4df1e4a9f111d6ae211816916?reference=9.

FDA. 2011. Coumadin (warfarin). https://wwwaccessdatafdagov/drugsatfda_docs/label/2011/009218s107lblpdf.

FDA. 2014. Soriatane (acitretin). https://wwwaccessdatafdagov/drugsatfda_docs/label/2014/019821s024lblpdf.

FDA. 2017. Doxycycline use by pregnant and lactating women. https://wwwfdagov/drugs/bioterrorism-and-drug-preparedness/doxycycline-use-pregnant-and-lactating-women.

FDA. 2019. Amoxicillin use by pregnant and lactating women exposed to anthrax. https://wwwfdagov/drugs/bioterrorism-and-drug-preparedness/amoxicillin-use-pregnant-and-lactating-women-exposed-anthrax.

Ferring. 2008. Endometrin® (progesterone). https://wwwaccessdatafdagov/drugsatfda_docs/label/2008/022057s001lblpdf.

Feutz AC, De Geyter C. 2019. Accuracy, discriminative properties and reliability of a human esc-based in vitro toxicity assay to distinguish teratogens responsible for neural tube defects. Arch Toxicol. 93(8):2375-2384.

Genschow E, Spielmann H, Scholz G, Pohl I, Seiler A, Clemann N, Bremer S, Becker K. 2004. Validation of the embryonic stem cell test in the international ecvam validation study on three in vitro embryotoxicity tests. Alternatives to laboratory animals: ATLA. 32(3):209-244.

GlaxoSmithKline. 2003. Myleran® (busulfan) https://wwwaccessdatafdagov/drugsatfda_docs/label/2003/09386slr023_myleran_lblpdf.

GlaxoSmithKline. 2014. Tafinlar (dabrafenib) capsules, for oral use. https://wwwaccessdatafdagov/drugsatfda_docs/label/2014/202806s002lblpdf.

Güveli BT, Rosti R, Güzeltaş A, Tuna EB, Ataklı D, Sencer S, Yekeler E, Kayserili H, Dirican A, Bebek N et al. 2017. Teratogenicity of antiepileptic drugs. Clinical psychopharmacology and neuroscience : the official scientific journal of the Korean College of Neuropsychopharmacology. 15(1):19-27.

Gwer SO, Onyango KO. 2018. Prevalence and incidence of congenital anomalies amongst babies born to women with sickle cell disease and exposed to hydroxyurea during pregnancy: A systematic review protocol. JBI Evidence Synthesis. 16(5):1135-1140.

Hochreiter S, Clevert D-A, Obermayer K. 2006. A new summarization method for affymetrix probe level data. Bioinformatics. 22(8):943-949.

Hyoun SC, Običan SG, Scialli AR. 2012. Teratogen update: Methotrexate. Birth defects research Part A, Clinical and molecular teratology. 94(4):187-207.

ICH. 2020. Ich harmonized guideline detection of reproductive and developmental toxicity for human pharmaceuticals s5(r3).

Kahn M. 2014. Can we safely target the wnt pathway? Nat Rev Drug Discov. 13(7):513-532.

Kam RK, Deng Y, Chen Y, Zhao H. 2012. Retinoic acid synthesis and functions in early embryonic development. Cell Biosci. 2(1):11.

King Pharmaceuticals I. 2012. Bicillin® c-r (penicillin g benzathine and penicillin g procaine injectable suspension) https://wwwaccessdatafdagov/drugsatfda_docs/label/2012/050138s234lblpdf.

Khanin R, Wit E. 2006. How scale-free are biological networks. Journal of Computational Biology. 13(3):810-818.

Kozma C, Ramasethu J. 2011. Methotrexate and misoprostol teratogenicity: Further expansion of the clinical manifestations. American journal of medical genetics Part A. 155a (7):1723-1728.

Langfelder P, Horvath S. 2008. Wgcna: An r package for weighted correlation network analysis. BMC Bioinformatics. 9(1):559.

Lammer EJ, Chen DT, Hoar RM, Agnish ND, Benke PJ, Braun JT, Curry CJ, Fernhoff PM, Grix AW, Jr., Lott IT et al. 1985. Retinoic acid embryopathy. The New England journal of medicine. 313(14):837-841.

Li R, Kimoto E, Niosi M, Tess DA, Lin J, Tremaine LM, Di L. 2018. A study on pharmacokinetics of bosentan with systems modeling, part 2: Prospectively predicting systemic and liver exposure in healthy subjects. Drug Metabolism and Disposition.dmd.117.078808.

Lipinski RJ, Hutson PR, Hannam PW, Nydza RJ, Washington IM, Moore RW, Girdaukas GG, Peterson RE, Bushman W. 2008. Dose- and route-dependent teratogenicity, toxicity, and pharmacokinetic profiles of the hedgehog signaling antagonist cyclopamine in the mouse. Toxicol Sci. 104(1):189-197.

Ljosa V, Caie PD, ter Horst R, Sokolnicki KL, Jenkins EL, Daya S, Roberts ME, Jones TR, Singh S, Genovesio A et al. 2013. Comparison of methods for image-based profiling of cellular morphological responses to small-molecule treatment. Journal of Biomolecular Screening. 18(10):1321-1329.

Lloyd KA. 2013. A scientific review: Mechanisms of valproate-mediated teratogenesis. Bioscience Horizons: The International Journal of Student Research. 6.

Martin MP, Bridgforth E, McGanity WJ, Darby WJ. 1957. The vanderbilt cooperative study of maternal and infant nutrition. X. Ascorbic acid. The Journal of nutrition. 62(2):201-224.

Matyskiela ME, Couto S, Zheng X, Lu G, Hui J, Stamp K, Drew C, Ren Y, Wang M, Carpenter A et al. 2018. Sall4 mediates teratogenicity as a thalidomide-dependent cereblon substrate. Nat Chem Biol. 14(10):981-987.

MaynePharma. 2008. Doryx® (doxycycline hyclate). https://wwwaccessdatafdagov/drugsatfda_docs/label/2008/050795s005lblpdf.

Mercer BM, Abdelrahim A, Moore RM, Novak J, Kumar D, Mansour JM, Perez-Fournier M, Milluzzi CJ, Moore JJ. 2010. The impact of vitamin c supplementation in pregnancy and in vitro upon fetal membrane strength and remodeling. Reproductive sciences (Thousand Oaks, Calif). 17(7):685-695.

Merck & Co. I. 2019. Decadron® (dexamethasone tablets, usp) https://wwwaccessdatafdagov/drugsatfda_docs/label/2019/011664s064lblpdf.

Milunsky A, Graef JW, Gaynor MF. 1968. Methotrexate-induced congenital malformations: With a review of the literature. The Journal of Pediatrics. 72(6):790-795.

Montoya-Eguía SL, Garza-Ocañas L, Badillo-Castañeda CT, Tamez-de la O E, Zanatta-Calderón T, Gómez-Meza MV, Garza-Ulloa H. 2015. Comparative pharmacokinetic study among 3 metformin formulations in healthy mexican volunteers: A single-dose, randomized, open-label, 3-period crossover study. Current Therapeutic Research. 77:18-23.

Moreno J, Belmont A, Jaimes O, Santos JA, López G, Campos MG, Amancio O, Pérez P, Heinze G. 2004. Pharmacokinetic study of carbamazepine and its carbamazepine 10,11-epoxide metabolite in a group of female epileptic patients under chronic treatment. Archives of medical research. 35(2):168-171.

Morris CA, Duparc S, Borghini-Fuhrer I, Jung D, Shin C-S, Fleckenstein L. 2011. Review of the clinical pharmacokinetics of artesunate and its active metabolite dihydroartemisinin following intravenous, intramuscular, oral or rectal administration. Malaria Journal. 10(1):263.

Muanda FT, Sheehy O, Bérard A. 2017. Use of antibiotics during pregnancy and the risk of major congenital malformations: A population-based cohort study. British journal of clinical pharmacology. 83(11):2557-2571.

Nahum GG, Uhl K, Kennedy DL. 2006. Antibiotic use in pregnancy and lactation: What is and is not known about teratogenic and toxic risks. Obstetrics and gynecology. 107(5):1120-1138.

Nau H. 1993a. Embryotoxicity and teratogenicity of topical retinoic acid. Skin Pharmacology and Physiology. 6(suppl 1)(Suppl. 1):35-44.

Nau H. 1993b. Embryotoxicity and teratogenicity of topical retinoic acid. Skin pharmacology: the official journal of the Skin Pharmacology Society. 6 Suppl 1:35-44.

Newton PN, Chaulet JF, Brockman A, Chierakul W, Dondorp A, Ruangveerayuth R, Looareesuwan S, Mounier C, White NJ. 2005. Pharmacokinetics of oral doxycycline during combination treatment of severe falciparum malaria. Antimicrobial agents and chemotherapy. 49(4):1622-1625.

Novartis. 2012. Gleevec (imatinib mesylate). https://wwwaccessdatafdagov/drugsatfda_docs/label/2012/021588s035lblpdf.

Paulson RJ, Collins MG, Yankov VI. 2014. Progesterone pharmacokinetics and pharmacodynamics with 3 dosages and 2 regimens of an effervescent micronized progesterone vaginal insert. The Journal of Clinical Endocrinology & Metabolism. 99(11):4241-4249.

Pfizer. 2007. Motrin®ibuprofen tablets, usp https://wwwaccessdatafdagov/drugsatfda_docs/label/2007/017463s105lblpdf.

Pfizer Do. 2002. Zyrtec®(cetirizine hydrochloride). https://wwwaccessdatafdagov/drugsatfda_docs/label/2002/19835s15,%2020346s8lblpdf.

Piñeyro-Garza E, Gómez-Silva M, Gamino Peña ME, Palmer J, Berber A. 2015. Isotretinoin conundrum: A randomized, openlabel, crossover study in mexico to evaluate the bioavailability and bioequivalence of three pharmaceutical preparations of isotretinoin in healthy participants. International journal of clinical pharmacology and therapeutics. 53(10):897-904.

Powell H, Ekert H. 1971. Methotrexate-induced congenital malformations. Medical Journal of Australia. 2(21):1076-1077.

Qu M, Xiong L, Lyu Y, Zhang X, Shen J, Guan J, Chai P, Lin Z, Nie B, Li C et al. 2021. Establishment of intestinal organoid cultures modeling injury-associated epithelial regeneration. Cell Research. 31(3):259-271.

Roche. 2010. Accutane®(isotretinoin capsules) https://wwwaccessdatafdagov/drugsatfda_docs/label/2010/018662s060lblpdf.

Rousselot P, Boucher Sp, Etienne G, Nicolini FE, Chauzit E, Makhoul PC, Coiteux Vr, Guerci As, Gardembas M, Legros L et al. 2010. Pharmacokinetics of dasatinib as a first line therapy in newly diagnosed cml patients (optim dasatinib trial): Correlation with safety and response. Blood. 116(21):3432-3432.

Rumbold A, Ota E, Nagata C, Shahrook S, Crowther CA. 2015. Vitamin c supplementation in pregnancy. Cochrane Database of Systematic Reviews. (9).

Sakata T, Chen JK. 2011. Chemical 'jekyll and hyde's: Small-molecule inhibitors of developmental signaling pathways. Chem Soc Rev. 40(8):4318-4331.

Sergejew T. 2015. Evaluation of a human embryonic stem cell-based screening assay for the identification of teratogenic pharmaceuticals. THESIS FOR MASTER OF ADVANCED STUDIES IN TOXICOLOGY.

Shepard TH, Lemire RJ. 2004. Catalog of teratogenic agents. JHU Press.

Shimozono S, Iimura T, Kitaguchi T, Higashijima S-i, Miyawaki A. 2013. Visualization of an endogenous retinoic acid gradient across embryonic development. Nature. 496:363.

Stephenson CM, Levin RD, Spector T, Lis CG. 2013. Phase i clinical trial to evaluate the safety, tolerability, and pharmacokinetics of high-dose intravenous ascorbic acid in patients with advanced cancer. Cancer chemotherapy and pharmacology. 72(1):139-146.

TevaPharmaceuticals. 2014. Cyproheptadine hydrochloride tablets usp. https://wwwiodinecom/drug/periactin/fda-package-insert.

Thauvin-Robinet C, Maingueneau C, Robert E, Elefant E, Guy H, Caillot D, Casasnovas RO, Douvier S, Nivelon-Chevallier A. 2001. Exposure to hydroxyurea during pregnancy: A case series. Leukemia. 15(8):1309-1311.

Tomson T, Battino D, Perucca E. 2019. Teratogenicity of antiepileptic drugs. Current opinion in neurology. 32(2):246-252.

Tsankov AM, Akopian V, Pop R, Chetty S, Gifford CA, Daheron L, Tsankova NM, Meissner A. 2015a. A qpcr scorecard quantifies the differentiation potential of human pluripotent stem cells. Nat Biotechnol. 33(11):1182-1192.

Tsankov AM, Gu H, Akopian V, Ziller MJ, Donaghey J, Amit I, Gnirke A, Meissner A. 2015b. Transcription factor binding dynamics during human es cell differentiation. Nature. 518(7539):344-349.

Vargesson N. 2019. The teratogenic effects of thalidomide on limbs. Journal of Hand Surgery (European Volume). 44(1):88-95.

von Mehren M, Widmer N. 2011. Correlations between imatinib pharmacokinetics, pharmacodynamics, adherence, and clinical response in advanced metastatic gastrointestinal stromal tumor (gist): An emerging role for drug blood level testing? Cancer treatment reviews. 37(4):291-299.

Walker G, Mandagere A, Dufton C, Venitz J. 2009. The pharmacokinetics and pharmacodynamics of warfarin in combination with ambrisentan in healthy volunteers. British journal of clinical pharmacology. 67(5):527-534.

Warkus ELL, Marikawa Y. 2018. Fluoxetine inhibits canonical wnt signaling to impair embryoid body morphogenesis: Potential teratogenic mechanisms of a commonly used antidepressant. Toxicological Sciences. 165(2):372-388.

Wasz-Höckert O, Nummi S, Vuopala S, Jáurvinen PA. 1970. Transplacental passage of azidocillin, ampicillin and penicillin g during early and late pregnancy. Scandinavian Journal of Infectious Diseases. 2(2):125-130.

Whitlow S, Bürgin H, Clemann N. 2007. The embryonic stem cell test for the early selection of pharmaceutical compounds. ALTEX-Alternatives to animal experimentation. 24(1):3-7.

Worley KE, Rico-Varela J, Ho D, Wan LQ. 2018. Teratogen screening with human pluripotent stem cells. Integr Biol (Camb). 10(9):491-501.

Young DW, Bender A, Hoyt J, McWhinnie E, Chirn G-W, Tao CY, Tallarico JA, Labow M, Jenkins JL, Mitchison TJ et al. 2008. Integrating high-content screening and ligand-target prediction to identify mechanism of action. Nature Chemical Biology. 4(1):59-68.
